# Intracellular Protein Editing to Enable Incorporation of Non-Canonical Residues into Endogenous Proteins

**DOI:** 10.1101/2024.07.08.602493

**Authors:** Jenna N. Beyer, Yevgeniy V. Serebrenik, Kaitlyn Toy, Mohd. Altaf Najar, Nicole R. Raniszewski, Ophir Shalem, George M. Burslem

## Abstract

The ability to study proteins in a cellular context is crucial to our understanding of biology. Here, we report a new technology for “intracellular protein editing”, drawing from intein-mediated protein splicing, genetic code expansion, and endogenous protein tagging. This protein editing approach enables us to rapidly and site specifically install residues and chemical handles into a protein of interest. We demonstrate the power of this protein editing platform to edit cellular proteins, inserting epitope peptides, protein-specific sequences, and non-canonical amino acids (ncAAs). Importantly, we employ an endogenous tagging approach to apply our protein editing technology to endogenous proteins with minimal perturbation. We anticipate that the protein editing technology presented here will be applied to a diverse set of problems, enabling novel experiments in live mammalian cells and therefore provide unique biological insights.

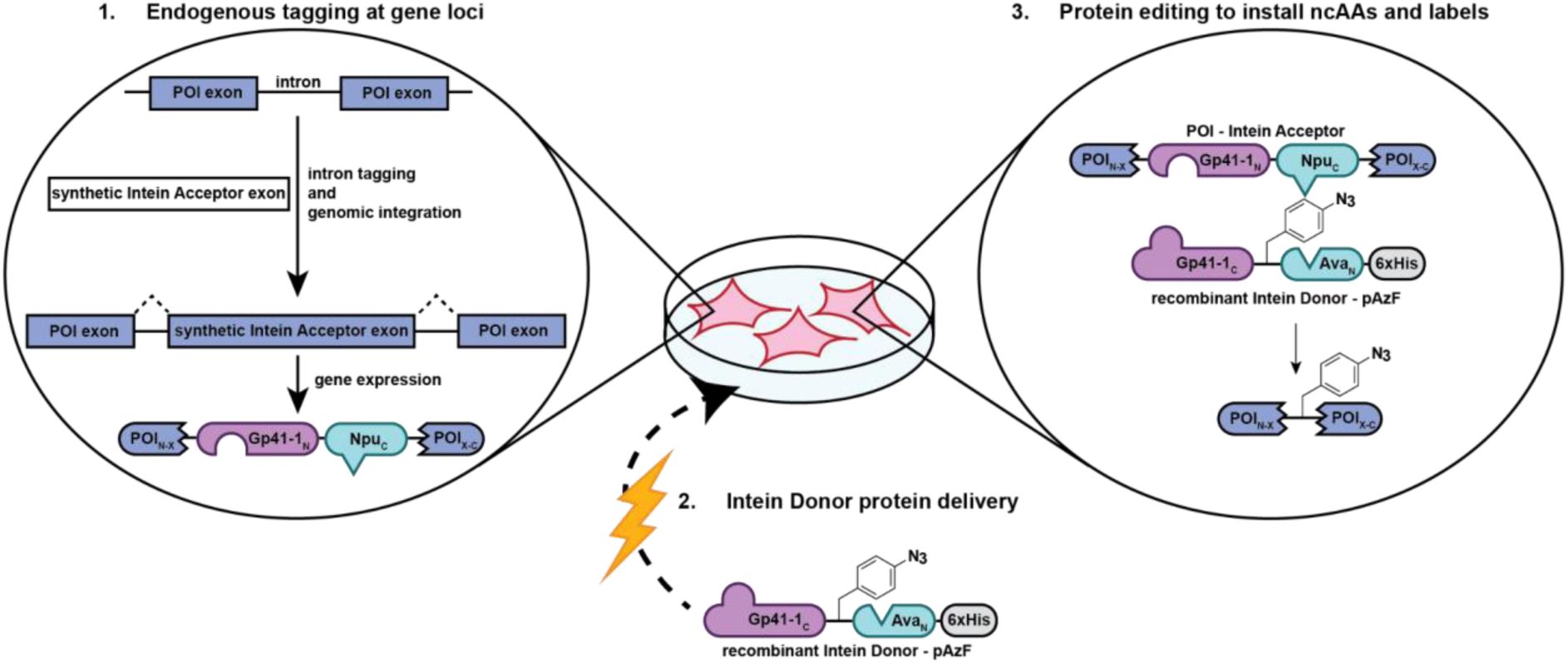

## Introduction

New technologies are imperative for the advancement of biomedical science, with novel strategies constantly being developed to enable researchers to better dissect and understand cellular interactions, pathways, and activities with precise detail. A clear example is the development of Cas9 gene editing, which has transformed biomedical research^1^. Gene editing strategies now routinely enable the precise manipulation of genes at endogenous loci, including the addition of epitope tags, fluorescent proteins and ligandable domains to gene products^2–7^. This advance has greatly improved experimental access to handles on proteins in mammalian cells, where proteins can be studied in a biologically relevant context. However, these gene editing approaches are limited to standard genetically encodable elements. In contrast, the conjugation of small molecules to proteins can offer similar and expanded functionalities (fluorescence and visualization, isolation, photocross-linking, and others)^8^ to genetically encoded tags but with minimal size and as such, less potential disruption. Currently, it is difficult for the cell biology and biochemistry fields to access the advantages of these small molecules-protein conjugates inside mammalian cells, leaving a gap for the development of new technologies.

Adding a tag to a protein of interest is the current benchmark for cell biology and biochemistry experiments in mammalian cells, in order to gain experimental access to the protein to be studied. At a broad level, tags enable a protein of interest to be more easily visualized in cells, identified as part of an interaction network, or otherwise tracked during experiments^9^. While incredibly useful and widely used, there are some caveats to consider when adding a tag to a protein of interest. Traditional protein tags range from short peptides, like a FLAG epitope tag (1.2 kDa), to a relatively large protein like Green Fluorescent Protein (GFP, 28 kDa). The tag itself, as well as the location of the tag^10,11^, can impact key characteristics of the fusion protein, such as its localization or protein-protein interactions^12–16^. In addition, using small molecules as tags can enable new experiments and reduce the size requirement of traditional tags. Methods such as SNAP-tag^17^ or HaloTag^18^ have recently pioneered the addition of small molecules to proteins in mammalian cells, but these approaches still require lengthy peptide or protein domain additions^8^. Together, these considerations demonstrate that there is no universal protein tagging strategy – instead, new and diverse tagging technologies are required to improve access to proteins and preserve their characteristics.

Another consideration when investigating a protein in mammalian cells is the current standard of investigating biological phenomena at a “steady-state,” with very little temporal resolution. It is known that signal transduction, for example, occurs rapidly, mostly on the minute to hours timescale^19–21^. However, routine cell biology experiments, such as overexpression, knockdown/knockout, or mutagenesis, largely lack the ability to study changes to proteins on such short timescales in cells. For example, in order to study a signaling network, researchers might induce long-term changes to signaling in mammalian cells by serum-starving cells, overexpressing a gene, or mutating a key residue to a post-translational modification (PTM) mimic, which are experiments that might change the signaling network for hours to days, in direct conflict with the dynamic timescales of biochemistry in the cell. Similarly, the constant and high level of expression of genes when transiently expressed can lead to confounding global changes throughout the cell, particularly when working with oncogenes^22^. This mismatch in kinetics, between what is currently experimentally accessible in mammalian cells and that of native biochemistry in cells, is known but cannot be easily remedied in routine cell biology experiments at present.

Therefore, in response to these gaps in the field, we envisioned a technology that could enable the rapid and modular editing of polypeptide primary sequences, including the addition of useful non-proteinogenic functional groups to both exogenously expressed and endogenous proteins, inside living mammalian cells (Fig. 1a). To realize this, we have developed a protein editing approach which employs split intein-mediated trans-protein splicing (TPS) to edit primary protein sequences. Importantly, we demonstrate that we can incorporate this editable sequence into endogenous proteins by genome editing. Furthermore, we can deliver exogenous protein, produced by genetic code expansion (GCE), and labeled by chemical biology approaches, into cells where it then undergoes editing to incorporate non-canonical functionality into endogenous proteins within minutes.

**Figure 1.**
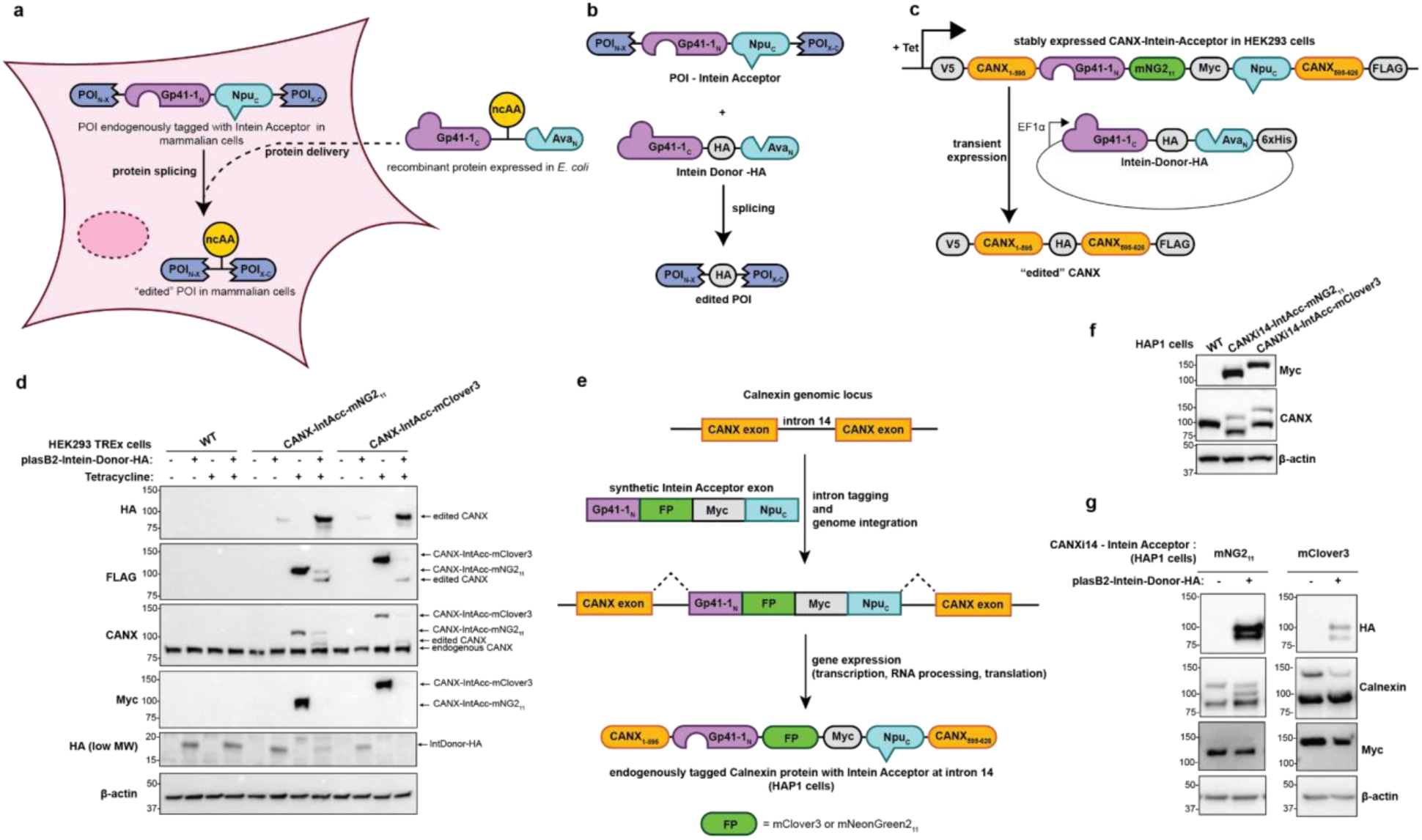
- A tandem split intein approach enables peptide swapping in proteins in mammalian cells. a, A general schematic for protein editing in mammalian cells using recombinant protein. **b,** Schematic for the arrangement of 2 orthogonal split inteins to enable dual splicing and the insertion of an Intein Donor peptide into a Protein of Interest (POI) – Intein Acceptor construct. **c,** Schematic of a FLAG-tagged calnexin with an Intein Acceptor construct, containing Myc and either mClover3 or mNG211 at the site corresponding to intron 14 (after exon 14) under Tetracycline control in stable HEK293 TREx FlpIn cell lines, where Intein Donor – HA is transiently expressed and results in an edited calnexin. **d,** Western blots resulting from the experiment in (c) show edited calnexin species now containing an HA tag (∼100 kDa). **e,** Endogenous proteins (in this case, calnexin) can be tagged with an Intein Acceptor construct using an intron tagging approach. **f,** Western blot of HAP1 cells with endogenous calnexin - Intein Acceptor (containing Myc epitope tag and either mClover3 or mNG211) inserted at intron 14 (between exons 14 and 15)**. g,** Transient expression of Intein Donor bearing HA in the endogenously tagged HAP1 cell lines results in the installation of an HA epitope tag into calnexin.

## Results

### Selection of Split Inteins and Configuration for Tandem Protein Trans-Splicing

To establish a platform for the modification of protein primary sequence in mammalian cells, we drew inspiration from previously established split intein technologies, pioneered by the Muir laboratory^23^. Several studies have shown that two split intein pairs can be used in tandem, arranged in such a way that the middle portion between the two orthogonal pairs can be exchanged^24,25^. Further, a recent publication from the Pless group demonstrated that proteins in mammalian cells could be modified in this way, but their efforts rely on synthetic peptides, which can be time and resource intensive to prepare^24^. We reasoned that a similar approach could be amenable to recombinant proteins and coupled to bacterial GCE for the facile incorporation of non-canonical amino acids (ncAAs) (Fig. 1a).

For our platform, we selected two well-characterized split intein pairs: Gp41-1 and AvaN:NpuC. Gp41-1 is the fastest characterized split intein to date, with a splicing half-life of ∼ 5 seconds under physiological conditions^26^. AvaN:NpuC is a recent derivative of the robust and widely used DnaE Npu split intein pair, also with a relatively fast splicing half-life of ∼1 minute *in vitro*^27,28^. Both split intein pairs possess different extein residues, required for splicing, that flank the splice site (Fig. S1a). When the various *N-* and *C-* terminal domains of the split inteins are separated onto two constructs, termed the “Intein Acceptor’’ and “Intein Donor”, and arranged as shown in Fig 1b, the residues between the inteins on the Donor can be inserted into an Acceptor sequence. Upon splicing, the four split intein domains are excised from the protein construct, leaving a nearly traceless “edit” to a protein’s primary sequence (Fig. S1b).

### A Model System for Protein Editing via Transient Expression

To test the trans-protein splicing concept, we constructed a model system using fluorescent proteins expressed from plasmids, where we appended an Intein Acceptor construct to the *C*-terminus of mClover3. The corresponding Intein Donor contained mRuby and an HA epitope tag, or an HA tag alone. With this configuration, if dual splicing is successful, the payload from the Intein Donor would be appended onto the *C*-terminus of mClover3 (Fig. S1c). We transiently expressed these two Intein Donor constructs with the mClover3 Intein Acceptor construct in human immortalized HEK293T cells and observed the expected 58 kDa (mClover3 + mRuby3-HA) or 33 kDa (mClover3 + HA) product bands by Western blotting (Fig. S1d).

Next, to test this on a functional human protein, we selected calnexin as a model for our system. Calnexin (CANX) is a chaperone protein that is located at the endoplasmic reticulum (ER) membrane, with a cytosolic *C*-terminal tail. Importantly, calnexin has been successfully endogenously and internally tagged in the past and has a distinct ER localization^2,29^. We designed a plasmid for transient expression in mammalian cells, starting with FLAG-tagged calnexin and inserting our Intein Acceptor construct after exon 14, which places the Intein Acceptor tag in the cytosolic *C*-terminal region. This “Intein Acceptor’’ construct contained the two intein domains, Gp41-1N and NpuC, with a fluorescent protein mClover3 and a Myc epitope tag situated between them, resulting in a 43 kDa addition (Fig. S1e). We additionally created a variant with mNeonGreen211 (mNG211)^30^, a single β-strand of a split GFP, in place of mClover3, to minimize the overall size of the Intein Acceptor insert to 18 kDa. Upon the transient expression of both Intein Acceptor-tagged calnexins in combination with the transient expression of an Intein Donor bearing an HA tag in HEK293T cells, we observe the spliced calnexin that now contains an internal HA epitope tag (Fig. S1f).

We then generated a stable HEK293 TREx FlpIn cell line from the calnexin - Intein Acceptor constructs containing mClover and mNG211, with the modified calnexins under Tetracycline-inducible expression (Fig. S1g,h). Upon transiently expressing an Intein Donor bearing an HA tag in these cell lines, we were able to replicate our previous results corresponding to the dual intein-mediated splicing, or “editing”, of a HA tag into an internal site in calnexin, as shown through the generation of a new FLAG- and calnexin-reactive species (Fig. 1c,d).

In tandem, we used a Cas9-based intron tagging approach^2,3^ in HAP1 cells to endogenously tag intron 14 of calnexin with the Intein Acceptor constructs containing mClover3 or mNG211, respectively, such that the Intein Acceptor sequence is included after exon 14 of calnexin (Fig. 1e). Flow cytometry and cell sorting was used to analyze and purify cell lines to homogeneity (Fig. S2a-c). These endogenously tagged cell lines were validated by Western blotting, where the higher molecular weight Intein Acceptor tagged calnexin is apparent by blotting for Myc or calnexin (Fig. 1f). Upon transiently expressing the intein donor containing an HA epitope tag in these cells, we again observe the ∼100 kDa spliced calnexin (Fig. 1g). Therefore, demonstrating that this approach allows us to edit proteins at endogenous expression levels, enabling more precise and more biologically relevant investigations of a protein or phenomenon of interest within the cellular milieu.

### Using Purified Recombinant Intein Donor for Protein Editing

Next, to show that we can install a minimal internal chemical modification on exogenously expressed or endogenous proteins in human cells, we aimed to convert this to a fully protein-based system by generating recombinant Intein Donor protein in *E. coli*. Split inteins are known to be unstable and disordered proteins that are prone to insolubility when expressed recombinantly^23,31–33^. To remedy this, we added an *N*-terminal SUMO solubility tag to our Intein Donor construct bearing an HA tag. After significant optimization, we were able to purify this Intein Donor in sufficient yield (∼30 mg of pure protein per liter of culture) (Fig. S3a,b).

With purified Intein Donor protein in hand, we now turned our focus to delivering the protein into cells. Electroporation has been used to routinely deliver oligonucleotides and plasmids into mammalian cells, and has also been utilized to deliver protein into mammalian cells^34–36^. We electroporated Intein Donor protein bearing HA epitope tag into the HAP1 cell lines containing calnexin endogenously tagged with Intein Acceptor- mNG211. Only in the presence of both electroporation and donor protein, we observe a newly formed calnexin species that now includes an HA epitope tag, as well as the anticipated concurrent reduction in Myc signal (Fig. 2a,b). Importantly, no other conditions yielded the high molecular weight and HA reactive bands, which are diagnostic of successful splicing. Using a range of concentrations of Intein Donor protein in the electroporation solution with both HAP1 and the stable HEK293 TREx FlpIn cell lines, we found that the optimal concentration of delivered protein was 50 µM (Fig. S4a,b).

**Figure 2.**
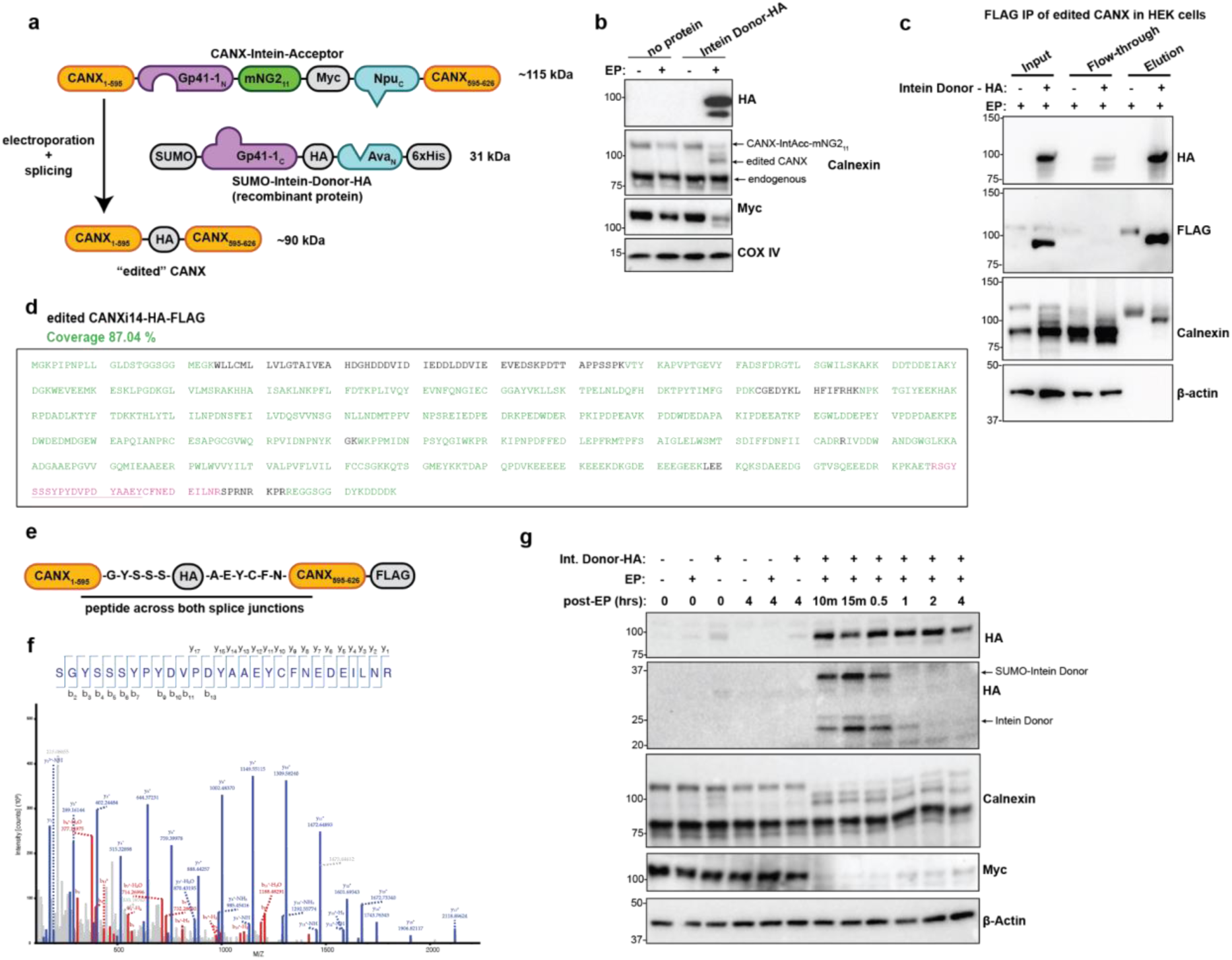
- Editing calnexin in mammalian cells using recombinant protein as a proof of concept. a, A schematic of protein editing approach using HAP1 cells with calnexin endogenously tagged with Intein Acceptor at intron 14 (after exon 14), via electroporation of recombinant Intein Donor protein bearing an HA tag as cargo. **b,** Western blot of resulting lysates from (a), showing edited calnexin now containing an HA epitope tag**. c,** FLAG resin pulldown of edited calnexin and un-edited calnexin-Intein Acceptor from the stable HEK293 TREx FlpIn cell lines, as shown by Western blotting. **d,** Sequence coverage of edited calnexin by mass spectrometry of the isolated edited calnexin from (c). The pink region is the peptide spanning the splice junctions and the underlined, pink region is the amino acid sequence that was edited into calnexin. Residues in green or pink correspond to peptides detected by MS. **e,** Schematic of a critical peptide that spans both split intein splicing sites in the edited calnexin product**. f,** The critical peptide from (e) identified by MS (in pink in d), with resulting spectra shown**. g,** A timecourse of calnexin splicing over 4 hours following electroporation of Intein Donor into HAP1 cells with endogenously tagged calnexin-Intein Acceptor-mNG211

In order to further confirm that both intein pairs were splicing, and that the expected edited calnexin was being generated, we employed mass spectrometry to validate the spliced product. Using our HEK293 TREx FlpIn stable cell line expressing FLAG-tagged calnexin- Intein Acceptor-mNG211, which provided sufficient material for testing the accuracy of protein editing by mass spectroscopy, we electroporated either Intein Donor bearing an HA tag or performed control electroporation (in the absence of Intein Donor) and incubated cells for 1 hour. We then lysed the cells, used anti-FLAG resin to isolate the FLAG-tagged calnexin species, and visualized the pulldown by Western blot (Fig. 2c). We next subjected the resulting protein to trypsin digestion and liquid chromatography tandem mass spectrometry (LC-MS/MS). Gratifyingly, we obtained coverage of the majority of the sequence for either the calnexin-Intein Acceptor species or the edited calnexin-HA species, depending on whether or not Intein Donor-HA was supplied prior to electroporation (Fig 2d, S5a). Critically, in the MS data for the edited calnexin-HA species, we observed the peptide corresponding to the two splice junctions, indicating that both inteins had spliced and formed the expected product (Fig. 2e,f). We also performed an interactome analysis using the edited calnexin-HA and identified 70 interactors that were previously reported for calnexin and are broadly associated with the ER (Fig. S5b, Table S1). Upon applying gene ontology (GO) analysis to the proteins identified in our interactome analysis, we find that 26 of the proteins are associated with the ER compartment and 14 are associated with the unfolded protein response. These are key characteristics of the ER-localized, chaperone protein calnexin and suggest that incorporation of the Intein Acceptor and subsequent editing to include an HA tag maintains ER localization and function.

Interestingly, when comparing editing in HAP1 cells with CANX endogenously tagged with Intein Acceptor containing either mClover3 or mNG211, we noticed that the two Intein Acceptors displayed different molecular weight products (Ext. Data Fig. 1a). Upon further investigation, we determined that the calnexin – Intein Acceptor with mClover3 was only splicing via the Gp41-1 split intein pair, leaving an incompletely spliced product. We confirmed this by generating a set of recombinant Intein Donors that were catalytically inactive for Gp41-1, Ava:Npu, or both, and used these mutants to validate that the Intein Acceptor containing mClover3 was not engaging the Ava:Npu split intein pair (Ext. Data Fig. 1c,d). For further discussion of this incomplete splicing, see Extended Text and Ext. Data Figure 1. In all following experiments, we used the Intein Acceptor containing mNG211 to promote splicing by both split intein pairs, in order to obtain our desired product.

We next investigated the splicing rate of our system within cells. While Gp41-1 and Ava:Npu are known to be individually fast and robust inteins *in vitro*^25–27^, it was unclear whether the split inteins could splice as efficiently when in close proximity to each other in a *N* and *C*-terminal constrained configuration inside mammalian cells. To test the speed of splicing in calnexin endogenously tagged with Intein Acceptor-mNG211, we harvested the cells at various time points following electroporation of Intein Donor bearing HA epitope tag (Fig. 2g and S4d). Excitingly, we observed completely spliced CANX at 10 minutes post-electroporation, the earliest time point possible following washes. Similarly, we also observe a near-complete depletion of the calnexin-Intein Acceptor species at the initial timepoints, followed by the gradual increase in signal over time as this protein is re- synthesized. This experiment demonstrated that dual splicing is rapid in the cell, enabling us to generate new protein species with precise temporal resolution. Further, when delivering Intein Donor into wild-type HAP1 cells that lack any Intein Acceptor, the Intein Donor protein is turned over quickly in cells and is undetectable by Western blot after 2 hours (Fig. S4d).

### Applying protein editing to a set of diverse cellular proteins

Having established this technology using calnexin as a model protein, we now wanted to apply it to a set of proteins occupying distinct locations and roles in the cell. We selected cytoskeleton components β-actin and α-actinin-1, the kinase Chk1, and transcription factor c-Myc as model systems to demonstrate our protein editing method’s broad utility across diverse proteins in the cell.

We selected β-actin (ACTB) as a target due to its abundance, longevity, and distinct localization pattern in the cytosol^37^. We inserted the Intein Acceptor with mNG211 after exon 2 and after exon 5 of β-actin (referred to as ACTBi2 and ACTBi5-Intein Acceptor, respectively) and generated a Tet-inducible stable HEK293 TREx FlpIn cell line for each (Fig. S6a). As before, we electroporated Intein Donor bearing HA into both ACTB tagged cell lines, with splicing resulting in robust product ACTB from both (Figs. 3a-c, S6b). We also tested the speed of splicing using the cell line with Intein Acceptor added after exon 2 of ACTB. Excitingly, splicing in this system also proceeds rapidly, with completely spliced β-actin forming within 10 minutes of electroporation of the Intein Donor protein (Fig. 3d), providing further evidence of the temporal control available using protein editing.

**Figure 3.**
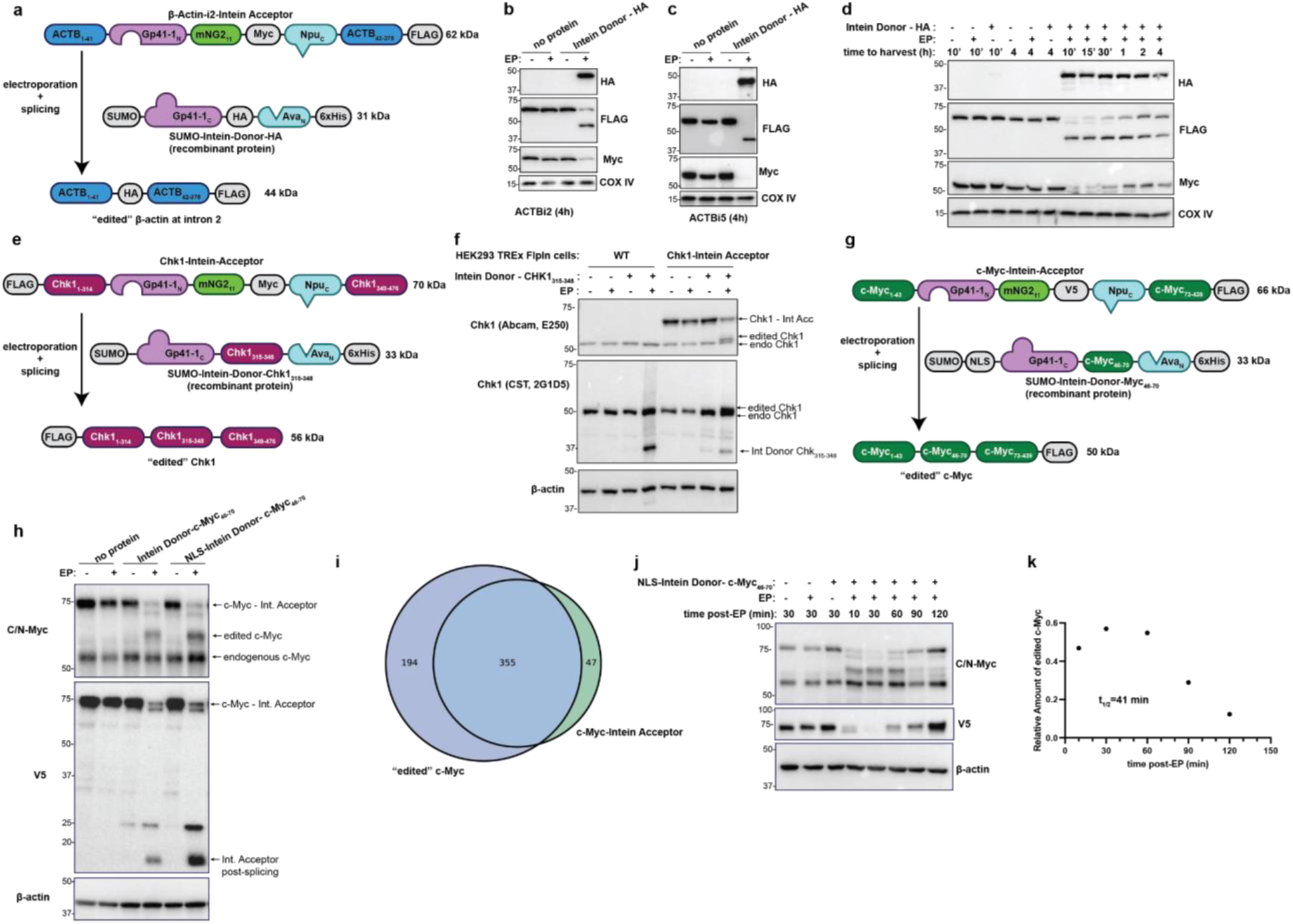
– Applying protein editing to a set of diverse proteins in the cell and demonstrating kinetic control. a, Schematic of protein editing approach in β-actin, using HEK293 TREx FlpIn cells stably expressing FLAG-tagged β-actin with Intein Acceptor at a site corresponding to intron 2, between exons 2 and 3. **b,** Western blot showing the editing of β-actin to now include an HA epitope tag at the site following exon 2**. c,** Western blot showing the editing of β-actin to now include an HA epitope tag at the site corresponding to intron 5. **d,** A timecourse of β-actin splicing following electroporation of Intein Donor bearing HA into HEK293 TREx FlpIn cells stably expressing FLAG-tagged β-actin with Intein Acceptor corresponding to after exon 2. Minutes are marked with a ‘ symbol. **e,** Schematic for protein editing approach with Chk1 using HEK293 TREx FlpIn cells stably expressing Chk1 with an Intein Acceptor. **f,** Western blots resulting from the experiment in (e), comparing the electroporation of Intein Donor – HA into either wild-type (WT) HEK293 TREx cells or the HEK293 TREx cells stably expressing Chk1-Intein Acceptor, showing edited Chk1 at 4 hours post-electroporation. **g,** Schematic for protein editing approach with c-Myc using HEK293 TREx cells stably expressing c-Myc with an Intein Acceptor containing mNG211 and a V5 tag**. h,** Western blots from the experiment shown in (g) at 1 hour following electroporation, using Intein Donor bearing c-Myc46-70 either with or without an NLS sequence, showing edited c-Myc formation at ∼60 kDa. **i,** Comparison of the interactomes of c-Myc -Intein Acceptor or edited c-Myc to the previously reported c-Myc interactome. **j,** Timecourse experiment following the editing of c-Myc-Intein Acceptor to include the c-Myc46-70 sequence, showing degradation of the edited c-Myc protein species over time**. k,** Quantification of edited c-Myc levels from (j) following electroporation. Band densitometry was used to calculate relative amount of edited c-Myc by normalizing edited c-Myc levels to endogenous c-Myc levels from the same blot.

To test protein splicing on another endogenous protein, we also used the intron tagging approach to tag α-actinin-1 (ACTN1), another component of the cytoskeleton. We introduced an Intein Acceptor with mNG211 between exons 18 and 19, resulting in a HEK293 cell line containing endogenously tagged ACTN1 (Fig. S6d). We were able to show that upon electroporation and addition of Intein Donor bearing HA, we could observe the formation of edited ACTN1 now containing an HA epitope tag (Fig. S6c,e), as an additional example of the utility of our method to edit endogenous proteins.

Next, we applied our protein editing approach to Chk1, a kinase essential for proper cell function. Kinases are a major component of signaling networks in cells, and studying them can be difficult due to the lack of both kinetic control and specific inhibitors, as well as the potential for signaling pathway adaptation or re-wiring^38,39^. Chk1 has both nuclear and cytosolic populations and is known to be phosphorylated by the kinase ATR in response to double-stranded DNA breaks, subsequently acting as a messenger to activate other proteins^40^. We designed this protein system in a slightly modified manner; instead of selecting an location between 2 exons and splicing in additional residues containing an epitope tag, we planned to replace a region with the Intein Acceptor and then edit the original sequence back into the protein. This approach, coupled with careful planning for the Intein Acceptor construct, can result in a near traceless protein editing event, with only very few mutations to accommodate the extein residues required for splicing. In this case, we inserted an Intein Acceptor containing a Myc epitope tag and mNG211 in place of Chk1 residues 315-348. This location enables the native sequence of Chk1 to be maintained as much as possible upon splicing, with only 2 mutations required for extein residues (Fig. S6f,g). We generated a stable cell line in HEK293 TREx FlpIn cells, such that this Chk1-Intein Acceptor was Tet-inducible (Fig. S6h). Correspondingly, we generated an Intein Donor protein that contains Chk1315 – 348 as cargo (Fig. S3c). Upon electroporating Intein Donor – Chk1315-348 into our CHK1-Intein Acceptor stable cell line, we observe the formation of a new Chk1 species by Western blot that corresponds to the anticipated edited Chk1 (Fig. 3e,f). In addition, we verified that splicing was proceeding rapidly in Chk1, with edited Chk1 present at 1-hour post-splicing (Fig. S6i). Excitingly, we demonstrate that our technology can be applied to kinases, which represent an extremely dynamic subset of signaling proteins in the cell, where studies could benefit from kinetic control over these proteins.

### Time-resolved modeling of protein stability in c-Myc

We next wanted to apply our protein editing technology to investigate protein homeostasis and degradation with high temporal resolution. To study protein homeostasis and turnover, a translation elongation inhibitor, cycloheximide, is routinely used. Unfortunately, cycloheximide is cytotoxic to cells over long periods of time and can introduce confounding variables due to the global inhibition of translation^41^, thus driving the development of new methods to study protein stability^42–45^. Due to the rapid nature of our protein editing approach, with the ability to produce edited protein in 10 minutes, we were curious if this method might be amenable to studying protein stability without inhibitors or isotopic labeling. As an example, we selected c-Myc, an oncogene and transcription factor that is commonly mis-regulated across cancers^46^. Under normal conditions, c-Myc has a very short half-life of only 20-40 minutes^47^. We were curious if we could recapitulate this known phenotype but with very precise temporal resolution, essentially releasing a new cohort of c-Myc via editing and subsequently tracking this protein’s lifetime following editing. Therefore, we generated a c-Myc-Intein Acceptor construct with a *C*-terminal FLAG tag, where the Intein Acceptor includes mNG211 and a V5 epitope tag. In this case, we designed the c-Myc – Intein Acceptor construct such that the Intein Acceptor replaces residues 46-70, in the Myc Box I domain, with the fully spliced c-Myc containing just 4 point mutations to accommodate the extein requirements of Gp41- 1 and Ava:Npu (Fig. S7a).

We first observed splicing by transient expression of the c-Myc-Intein-Acceptor and Intein Donor constructs (Fig. S7b) and subsequently generated a HEK293 TREx FlpIn stable cell line expressing the c-Myc – Intein Acceptor (Fig. S7c). We produced a recombinant Intein Donor that included c-Myc45-70 as the payload (Fig. S3d) with an added NLS sequence since c-Myc is localized to the nucleus. Upon electroporating an Intein Donor bearing an NLS and the c-Myc45-70 sequence into the stable c-Myc Intein Acceptor cells, we can detect a new protein species that is reactive to C/N-Myc antibody on a Western blot. This spliced c-Myc is concurrent with a loss of V5 signal from the Intein Acceptor, confirming splicing of both inteins (Fig. 3g,h). Interestingly, our stable cell line expressing c-Myc-Intein Acceptor-FLAG was not immunoreactive to FLAG on Western blots, but we were able to isolate both c-Myc -Intein Acceptor and edited c-Myc using a FLAG pulldown (Fig. S7d). We subjected this immunoprecipitated c-Myc-Intein Acceptor and edited c-Myc to mass spectrometry and were able to confirm via peptide sequence coverage that this species is the expected edited c-Myc product (Fig. S7e). We were also able to use FLAG Co-IP-MS to obtain interactomes for both the c-Myc-Intein Acceptor and edited c-Myc (Fig. 3i, Table S2&3). These interactomes were broadly similar, demonstrating that the addition of the Intein Acceptor tag prior to editing does not significantly change interactions to the protein of interest, at least in the context of our c-Myc configuration, further highlighting the non-perturbing nature of our method. Importantly, the other component of the c-Myc transcription factor heterodimer complex, Max^46^, was identified as an interactor in both datasets, showing that this critical interaction is maintained whether c-Myc is tagged with Intein Acceptor or edited to a near wild-type sequence.

With our c-Myc splicing configuration validated, we turned now to measuring protein half-life following splicing. We electroporated Intein Donor bearing an NLS and c-Myc45-70 into the c-Myc-Intein Acceptor cells and harvested cells at time points following electroporation (Fig. 3j). Upon quantifying this Western blot data, we were able to see rapid turnover of the spliced c-Myc as expected (t1/2 = 41 minutes), consistent with the previously reported half-life under cycloheximide conditions^47^ (Fig. 3k). This represents a novel method to study protein half-lives and stability without the use of global chemical perturbations and offers a temporally resolved vantage point into the regulation of proteins in a way that was previously inaccessible.

### Expanding protein editing to include unnatural amino acids and useful labels

Now, having laid the groundwork and explored the versatility of our protein editing technology, we wanted to expand the method to enable the use of functionality beyond that which can be encoded genetically. To this end, we used genetic code expansion (GCE) in *E. coli* to introduce a click chemistry handle as cargo into the recombinant Intein Donor (Fig. S8a), such that the click chemistry handle could be labeled and then installed into a protein of interest in mammalian cells via our protein editing technology. GCE enables the encoding on ncAAs by using an engineered tRNA/aminoacyl-tRNA synthetase (aaRS) pair that are orthogonal to the host cell’s translation machinery. The orthogonal aaRS loads an ncAA onto the orthogonal tRNA, which suppresses an amber stop codon (TAG) in the target gene, resulting in the addition of the ncAA to the nascent peptide chain^48^.

We initially selected the unnatural amino acid p-azido-phenylalanine (pAzF) as a proof of concept due to its functionality as a click chemistry handle and photo-crosslinker^49,50^. We were able to express and purify Intein-Donor-pAzF following the workflow already established for Intein Donor containing canonical amino acids (Fig. S8b,c). This minimal Intein Donor bearing pAzF contains the several required extein residues and pAzF, enabling a very small addition into the protein of interest. Following purification of Intein Donor - pAzF, we selected a set of useful and interesting small molecule labels, including biotin, several fluorophores (Fluor488, TAMRA, and Alexa Fluor 647), sortase compatible Gly-Gly-Gly peptide^51^, and the sugar GlcNAc, to conjugate onto the Intein Donor protein using various click chemistry reactions (Fig. 4a). In particular, we focused on biotin and the fluorophore tetramethylrhodamine (TAMRA) as useful labels, as biotin would enable pulldown experiments and TAMRA could be used for microscopy applications. To conjugate these small molecules to Intein Donor - pAzF on a preparative scale, we routinely utilized Copper-free strain promoted azide-alkyne cycloaddition (SPAAC) reactions and labeled the protein with biotin-DBCO and TAMRA-DBCO reagents as this had no impact on the solubility of the protein. Copper-mediated azide-alkyne cycloaddition (CuAAC) reactions on Intein Donor – pAzF resulted in some protein precipitation, though through the use of BTTAA ligand this can be minimized (Fig. S8d)^52^.

**Figure 4.**
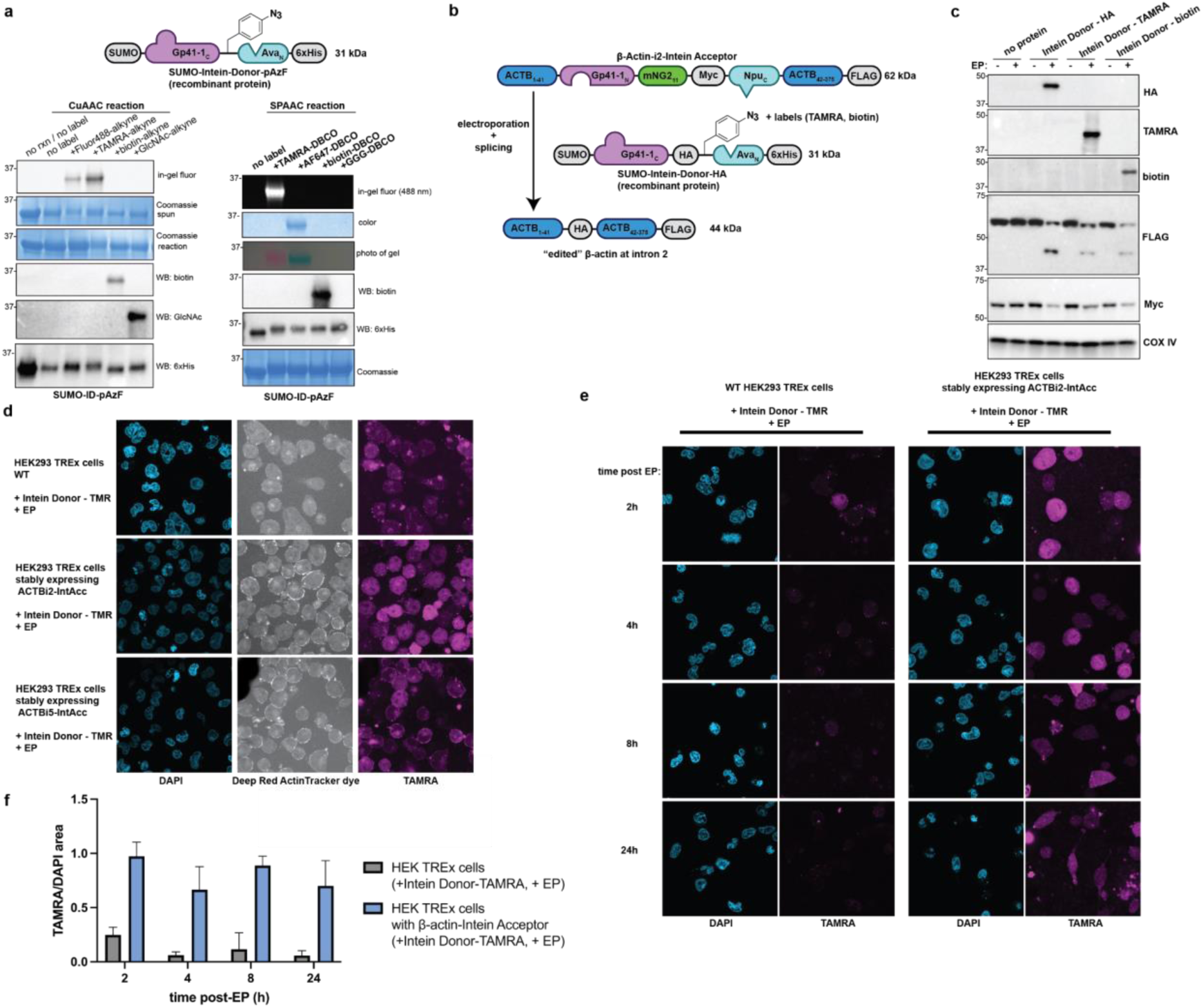
- Editing the ncAA p-azido-Phe (pAzF) and labels into calnexin and β-actin in stable cell lines. a, Labeling purified recombinant Intein Donor- pAzF with a variety of labels using CuAAC or Cu-free SPAAC reactions. **b,** Schematic for β-actin editing at intron 2, using recombinant Intein Donor bearing an HA epitope tag, or pAzF that was labeled with TAMRA or biotin. **c,** Western blot resulting from the experiment in (b), where the cargos from each Intein Donor have been edited into β-actin in stable HEK293 TREx FlpIn cell lines. **d,** Confocal microscopy images of Intein Donor – TAMRA electroporated into wild-type (WT) HEK293 TREx FlpIn cells or HEK293 TREx FlpIn cells stably expressing β-actin-Intein Acceptor at intron 2 or intron 5, with concurrent staining with DeepRed ActinTracker stain. **e,** A confocal microscopy timecourse with Intein Donor-TAMRA electroporated into wild-type (WT) HEK293 TREx FlpIn cells or HEK293 TREx Flpin cells stably expressing β-actin-Intein Acceptor at intron 2**. f,** Quantification of the microscopy timecourse in (e), where TAMRA area has been normalized to DAPI area for each field of view (n=3 or more per condition). Error bars represent one standard deviation.

We next wanted to test if Intein Donor protein bearing the broadly applicable labels TAMRA and biotin could be edited into proteins in mammalian cells via our approach. We first utilized the previously generated stable HEK293 TREx FlpIn cell lines with β-actin and calnexin containing Intein Acceptor. Using our stable cell line β-actin model with Intein Acceptor after exon 2, we electroporated Intein Donor bearing HA or Intein Donor bearing pAzF labeled with TAMRA or biotin into cells. Excitingly, upon Western blotting, we observe signals at ∼100 kDa for HA epitope tag, TAMRA, and biotin each in their respective donor conditions and only with electroporation. We also see FLAG signal corresponding to edited β-actin at the expected ∼44 kDa, and with the loss of Myc, this suggests that both split inteins are splicing (Fig. 4b,c). Having shown that our protein editing method enables the site-specific introduction of unnatural functionalities, such as TAMRA fluorophore and biotin, in living mammalian cells, we were excited by the numerous applications and experiments enabled by the post-translational and temporally-controlled installation of these useful functional groups.

Next, we wanted to use our ability to install fluorophores into β-actin to visualize the protein’s localization in cells, in hopes that our method would be accurate and low background when used with confocal microscopy. We electroporated Intein Donor bearing TAMRA into the HEK293 TREx FlpIn stable cells lines with ACTB tagged with Intein Acceptor after exon 2 or 5. After fixing and staining cells with an ActinTracker dye, we observed diffuse cytosolic distribution of TAMRA with some enrichment at the cell membrane, consistent with the ActinTracker dye (Fig. 4d). Further, we set up a time course experiment and fixed either wild-type or ACTB tagged with Intein Acceptor after exon 2 HEK293 FlpIn TREx cells at time points following electroporation of an Intein Donor bearing TAMRA (Fig. 4d,e). In the cells containing ACTB with Intein Acceptor after exon 2 and electroporated Intein Donor - TAMRA, we again observe TAMRA throughout cells and at the plasma membrane, while the TAMRA signal in wild-type HEK293 TREx FlpIn cells electroporated with Intein Donor- TAMRA is mainly localized to small puncta.

We note that while installing the TAMRA fluorophore between exon 5 and 6 produced an expected ACTB localization pattern, insertion at a different location between exon 2 and 3 resulted in a more diffuse localization (Fig. 4e). This is consistent with previous endogenous tagging of ACTB in these locations^2^, as well as other ACTB tagging studies^53^, and could suggest disruption to either the modified protein co-translationally or the assembly of the cytoskeleton structures. Gratifyingly, we see the TAMRA signal from the labeled ACTB persist over 24 hours, consistent with the long half-life of ACTB^54^, while the TAMRA signal from the wild-type cells diminishes rapidly to background levels by 4 hours post electroporation (Fig. 4e,f).

In addition, we completed a similar set of experiments in the HEK293 TREx FlpIn cell line stably expressing calnexin – Intein Acceptor. We demonstrated the installation of TAMRA and biotin into calnexin (Fig. S9a) and successfully isolated edited calnexin containing biotin from cell lysates using streptavidin resin (Fig. S9b). Further, we electroporated Intein Donor bearing TAMRA into either wild-type HEK293 TREx FlpIn or the stable HEK293 TREx FlpIn calnexin – Intein Acceptor cells and subsequently fixed cells for imaging at time points following initial delivery (Fig. S9c,d). Here, we observe an ER-localization pattern from the TAMRA signal, as we would expect from calnexin labeling. Importantly, the TAMRA signal from wild-type HEK293 TREx FlpIn cells electroporated with Intein Donor bearing TAMRA is diffuse throughout cells or in puncta, with signal dramatically decreasing over time, matching data obtained from the earlier β-actin imaging time course.

### Incorporation of Non-Canonical Residues in Endogenous Proteins

Lastly, we were eager to demonstrate the utility of our method for accessing and labeling endogenous proteins with non-disruptive and versatile chemical modification. Using the HAP1 cell line with endogenously tagged calnexin - Intein Acceptor, we electroporated a set of Intein Donors containing either an HA tag, or pAzF labeled with TAMRA or biotin as a cargo. Upon Western blotting for these respective cargos, we observe the molecular weight corresponding to the addition of the cargo to calnexin (Fig. 5a,b), providing evidence of the installation of non-canonical residues into endogenous proteins within living mammalian cells. In addition, we wanted to use the fluorescent Intein Donor - TAMRA to further validate electroporation as a suitable protein delivery method into cells. We used flow cytometry after electroporating Intein Donor - TAMRA into wild-type HAP1 cells to demonstrate that the fluorescent Intein Donor is being delivered into cells and that a significant population of cells remain healthy, using Sytox Blue as a marker for dead cells (Fig. S10a).

**Figure 5.**
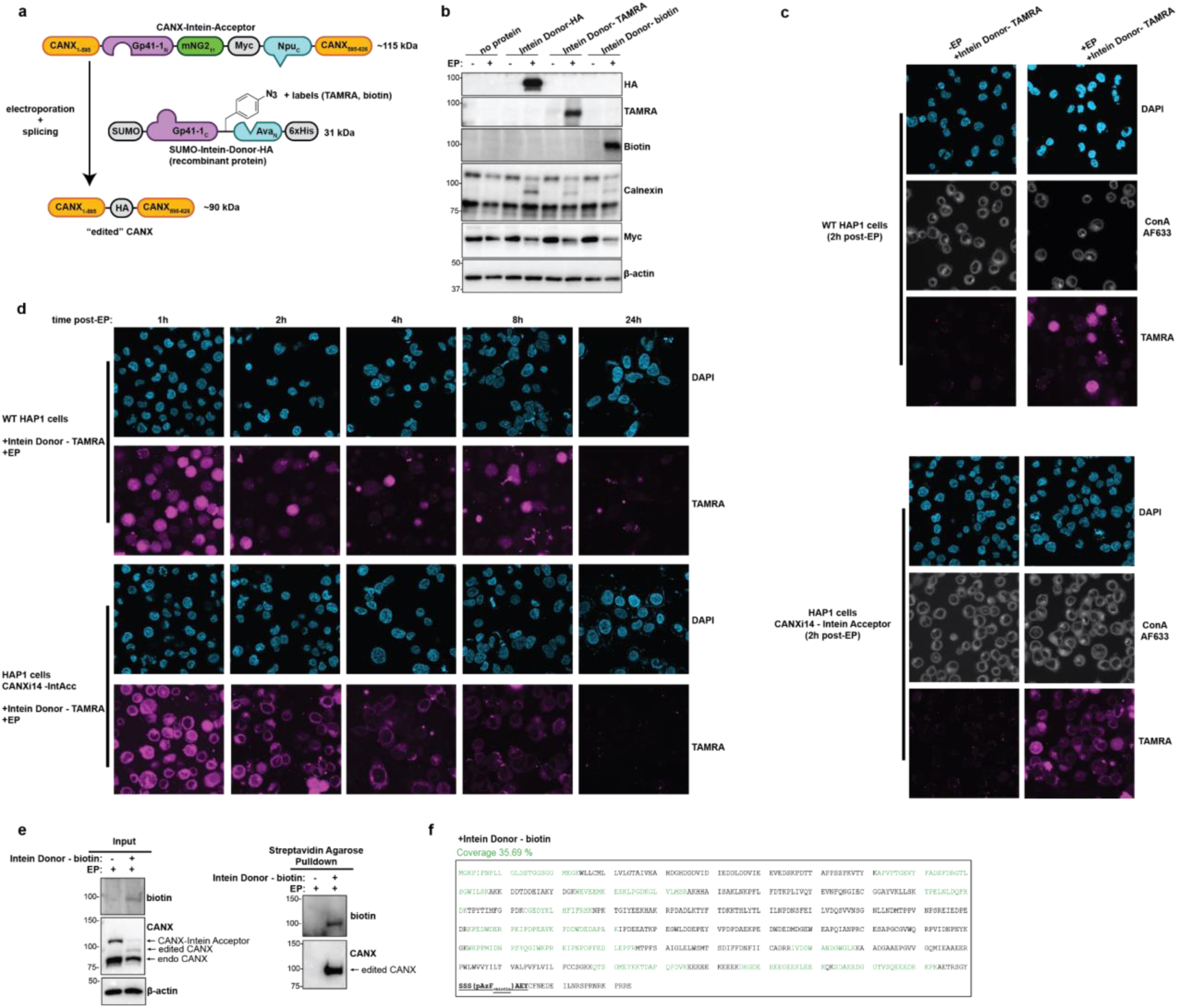
- Editing pAzF bearing useful labels into endogenous calnexin. **a,** Schematic for editing various cargos, either an HA tag or a label conjugated to pAzF by click chemistry, into endogenously tagged calnexin in HAP1 cells. **b,** Western blot of resulting lysates from (a) after 4 hours recovery post-electroporation. **c,** Confocal microscopy images comparing wild-type HAP1 cells and HAP cells with endogenous calnexin tagged with Intein Acceptor, both with and without electroporation of an Intein Donor bearing TAMRA. Cells were also stained with Concanavalin A – AlexaFluor 633 (ConA) to show ER localization within cells. **d,** Confocal microscopy images of either wild-type HAP1 cells or HAP1 cells with endogenous calnexin tagged with Intein Acceptor. Following electroporation of Intein Donor bearing TAMRA, cells were fixed at time points. **e,** Editing a biotin into endogenously tagged calnexin in HAP1 cells and isolating the edited protein by a pulldown with streptavidin agarose. **f,** Mass spectrometry analysis of the protein isolated in (e) reveals significant sequence coverage of calnexin. Residues in green correspond to peptides detected by MS. The bold, underlined residues correspond to the region that is edited into calnexin.

With the ability to install a fluorophore site specifically into calnexin, we next employed confocal microscopy to visualize the subcellular localization of our edited endogenous calnexin in HAP1 cells. As anticipated, the TAMRA fluorescence was localized to the ER as is consistent with calnexin labeling, as demonstrated by the concurrent staining with Concanavalin A (ConA), an ER marker^55^ (Fig. 5c). Importantly, the TAMRA signal was dependent on electroporation and presence of the TAMRA labeled Intein Donor protein. When we fixed HAP1 cells at various time points following electroporation, we could see immediate and specific TAMRA signal from the ER that was maintained over at least 8 hours. Contrastingly, when we electroporate TAMRA labeled Intein Donor into wild-type HAP1 cells, we initially observe diffuse fluorescence across the cell and within puncta, with TAMRA signal decreasing with time (Fig. 5d, S11a with ConA staining). These results are consistent with earlier immunoblotting experiments showing that unspliced Intein Donor is degraded in cells within 2-4 hours of delivery (Fig. 2g, S4c). These characteristics demonstrate that this protein editing method is both on target (with only ER-localized labeling in presence of tagged calnexin) and has a low background (with the degradation of unspliced TAMRA Intein Donor) that will be useful for the analysis of other proteins.

Finally, we sought to explore the potential of installing a biotin handle to enable pull-down experiments in the absence of antibodies. We were able to edit biotin into calnexin using the endogenously tagged HAP1 cells and isolate the protein using streptavidin agarose (Fig. 5e). We subjected this protein to LC-MS/MS and identified multiple diagnostic peptides for calnexin. This experimental set-up enables the isolation and analysis of proteins in mammalian cells without the use of antibodies, which may well prove useful for studies involving proteins that are incompatible with standard epitope tagging approaches or lack good antibodies.

## Discussion

Here, we have developed a novel platform for post-translationally editing the primary sequence of proteins in mammalian cells in such a way that we can include functionalities beyond the routinely genetically encodable and under endogenous expression conditions. Our method enables access to both exogenous and endogenous proteins, which are previously modified through the use of Cas9 gene editing approaches^2^. We have demonstrated this protein editing method across a set of diverse proteins with distinct functions and locations in the cell, including calnexin, β-actin, c-Myc, α-actinin-1, and Chk1. We have introduced useful functional groups such as fluorophores and biotin, thus enabling classic cell biology experiments such as microscopy to study protein localization or pulldowns to isolate a protein of interest, but without the bulky fusion protein tags or antibodies that are typically used. Additionally, we demonstrate the use of protein-specific Intein Donors to model a near wild-type sequence, enabling kinetic control over individual protein species, again under non-perturbing conditions.

Furthermore, this approach is extremely multiplexable, since one minimal Intein Donor protein bearing a pAzF could be labeled with diverse functional groups and edited into an almost unlimited number of different proteins. Because we use click chemistry to further functionalize the Intein Donor protein, one could add any of the widely available fluorophores, probes, peptides, nucleic acids, or other useful functional groups to the Intein Donor protein and use it for seemingly limitless downstream applications in mammalian cells. In addition, we have utilized GCE in *E. coli* to incorporate an ncAA into recombinant Intein Donor protein, with numerous other GCE-enabled ncAAs available to be incorporated into the Intein Donor protein. Further, we note that GCE in mammalian cells, while rapidly expanding, still faces some challenges: prematurely terminated protein by-products, overall burden of additional translation machinery, and background from readthrough of native stop codons^56^. Our method circumvents these challenges by using GCE in *E. coli* and labeling the purified Intein Donor protein *in vitro*, but ultimately yields site specific ncAA installation in mammalian cells.

Perhaps most importantly, our protein editing method is rapid, with high temporal resolution in which to probe cellular responses to the release of the edited protein species. We have shown that we can engage with this temporal resolution through protein stability experiments and by observing the temporal decay of fluorescence signal via confocal microscopy with calnexin and β-actin. In sum, we believe that this temporal resolution will prove powerful for studying native biochemistry in mammalian cells and will only continue to grow more crucial as the field better understands the temporal requirements of processes in the cell.

As with any method, our protein editing approach is not without limitations. Firstly, to be applied to endogenous protein, this method still relies on genome editing to insert the Intein Acceptor sequence which could potentially perturb the function of the targeted protein. Perhaps the biggest challenge is the delivery of the exogenous Intein Donor protein, which still proves challenging and relies on electroporation throughout this manuscript. However, recent advances in the field of protein delivery^57–59^ will likely enable new solutions and reduce these concerns. Further, the exploration of new ways to deliver Intein Donor protein is an active area in our labs.

In summary, here we present a new platform for protein editing, which enables the post-translational editing of the primary sequence of proteins in live mammalian cells. Our protein editing method is a rapid, temporally controllable, and highly multiplexable tool that can enable the access and modification of endogenous proteins in living mammalian cells. We believe that this technology will be widely applicable to many diverse proteins and will catalyze new fundamental understandings in cellular biochemistry.

## Acknowledgements

We would like to thank the Shalem and Burslem labs for extensive discussions related to this manuscript. This work was supported by the following grants: Discovering the Future Research Grant from the Office of the Vice Provost for Research at the University of Pennsylvania (to G.M.B.), a research grant from the Basser Center for BRCA research (to G.M.B.), DP2GM137416 from NIH/NIGMS, SAP#4100083086 from PA DoH and R03NS111447-01 from NINDS awarded to O.S., R35GM142505 from NIGMS awarded to G.M.B. F32CA239499 from NCI and K99AG075256 from NIA awarded to Y.V.S. JNB was supported by a NSF Graduate Research Fellowship. NRR was supported by the NIH Chemistry Biology Interface Training Grant (T32 GM133398). We thank the Penn CDB Microscopy Core (RRID SCR_022373) for the use of their instruments.

## Materials and Methods

### Molecular biology and cloning

All oligos were purchased from IDT. Gene fragments were purchased from Twist and IDT. PCRs were performed with 2xQ5 HotStart MasterMix (NEB). Plasmids were generated by Q5 Site-Directed Mutagenesis or Gibson Assembly (NEB). pDule2-pCNF was a gift from Ryan Mehl (Addgene plasmid # 85495; http://n2t.net/addgene:85495; RRID: Addgene_85495).

### General Cell Culture

HEK293T and HEK293 TREx FlpIn cells (ThermoFisher) were grown in DMEM (Gibco) supplemented with 10% FBS (Hyclone) and 1% Pen-Strep (Gibco). HEK293 TREx FlpIn cells were grown in the presence of Zeocin (100 µg/mL). HAP1 cells (Horizon Discovery) were grown in IMDM (Gibco) supplemented with 10% FBS (Hyclone) and 1% Pen-Strep (Gibco). Cells were split using DPBS (Gibco) and 0.25% Trypsin-EDTA (Gibco). Cells were passaged twice per week and used for experiments for 12-16 passages.

### Generation of endogenously tagged cell lines

Endogenously tagged cell lines were generated by homology-independent intron targeting, as reported previously^2^. Briefly, cells were transfected with 1) a plasmid expressing a gene intron-targeting sgRNA, 2) a plasmid encoding the DNA cassette for tagging, 3) a plasmid expressing an sgRNA to linearize the DNA cassette, and 4) a Cas9- expressing plasmid.

The DNA cassette encoding the tag consisted of mClover3-MYC or mNG211-MYC flanked by Gp41-1(N) and Npu(C). These sequences, in turn, are flanked by splice acceptor and donor sites to integrate the tag seamlessly into the coding sequence as a synthetic exon. Outside of the splice sites we added a constitutively-expressing blasticidin resistance gene to select for integration of the DNA cassette into the genome. Lastly, flanking all the previously mentioned DNA elements are sgRNA target sites for linearization of the DNA donor.

Following transfection and selection for genomic integrants with blasticidin, cells were selected for expression of fusion proteins by fluorescence-activated cell sorting (FACS). Prior to sorting mNG211-tagged cells, mNG21-10 was expressed by treatment with doxycycline (see “Generation of stable cell lines” for more detail).

### Generation of stable cell lines

HEK293 TREx FlpIn cells (ThermoFisher) were plated and subsequently transfected with a 9:1 ratio of pOG44:pcDNA5 vector containing the gene of interest. Several days after transfection, selection for positive clones was begun by adding 50 ug/mL of Hygromycin B to the media. Selection was continued for 2 weeks, with media changes every 2-3 days, until all mock cells had died. Cells were then allowed to expand without Hygromycin to yield a polyclonal population of cells. Cells were then tested with the addition of 1 µg/mL Tetracycline to confirm Tet-dependent expression of the gene of interest by immunoblotting.

For endogenous gene tagging using an mNG2(11)-based intein acceptor, we first established stable parental cell lines in HEK293 and HAP1 cells through lentiviral transduction to express mNG21-10 fused to the hygromycin resistance gene via a 2A peptide, driven by a doxycycline-inducible promoter (TRE-mNG21-10-2A-hygroR). Following transfection with the tagging reagents (see “Generation of endogenously tagged cell lines”), cells that stably expressed the tagged gene product were identified after treatment with 500 ng/μl doxycycline and subsequently selected by cell sorting. Alternatively, for tagging α-actinin-1 with an mNG211-based Intein Acceptor, we generated HEK293 cells constitutively expressing mNG21-10 fused to tdTomato via a 2A peptide.

### Expression and purification of Intein Donor constructs from E. coli

Expression plasmids were transformed into chemically competent BL21(DE3) cells (Invitrogen). For Intein Donor constructs with pAzF incorporated via GCE, BL21(DE3) cells were co-transformed with pET28 expression plasmid and pDule2-pCNF^50^. Transformed cells were plated onto LB plates with relevant antibiotics and allowed to grow at 37 °C overnight. 10-12 colonies were picked from the resulting plate and used to inoculate starter LB cultures with appropriate antibiotics, which were grown overnight at 37 °C. The larger expression culture was grown at 37 °C and 215 RPM until cells grew to an OD of 0.4-0.6, at which point protein expression was induced with the addition of 1 mM IPTG (and 1mM pAzF, if carrying out GCE). Typically, expressions were carried out at 37 °C overnight. Cells were pelleted by centrifugation at 12,000 rpm, 4°C for 30 minutes and then frozen at -80°C.

For purification, cell pellets were thawed on ice, and then resuspended in a lysis buffer containing 50 mM HEPES pH 7.5, 300 mM NaCl, 5% glycerol, and Protease Inhibitor (Pierce). Cell pellets were sonicated at 50% power for 10 minutes total (4s on, 2s off), and the lysate was subsequently cleared at 13,000 rpm for 30 min at 4 °C. Supernatant was retained, while a resolubilization buffer of 50 mM HEPES pH 7.5, 300 mM NaCl, 5% glycerol, and 8 M urea was added to the pellet and proteins resuspended. This re- solubilized pellet solution was left to rock overnight at 4 °C, and the following day, the solution was spun again at 13,000 rpm, 4 °C for 30 minutes. The resulting supernatant contains protein extracted from the earlier pellet and was used in subsequent purifications.

NiNTA resin was equilibrated with lysis buffer. The supernatant from the cell lysate and the pellet was applied to resin and allowed to rock for 1 hour or overnight at 4 °C. The column was washed with 2 washes of 50 mM HEPES pH 7.5, 300 mM NaCl, and 50 mM imidazole, and protein was subsequently eluted with 50 mM HEPES pH 7.5, 300 mM NaCl, and 300 mM imidazole. Protein was dialyzed against 50 mM HEPES pH 7.5 and 100 mM NaCl and then concentrated using a 10 kDa MWCO concentrator tube. Protein was then purified further by size exclusion chromatography on an S200 Sephadex column, and then concentrated again. Protein was snap frozen in liquid N2 and stored at -80 °C.

### Modification of Intein Donor protein by Click Chemistry

For Cu-free click chemistry, biotin-PEG4-DBCO (Vector Labs), TAMRA-DBCO (Vector Labs), and AlexaFluor647-DBCO (BroadPharm) were added to purified Intein Donor in a 5:1 molar ratio (label: protein) and protein solutions were left to rock overnight at room temperature.

For Cu-catalyzed azide-alkyne cycloaddition, 500 μM BTTAA (Vector Labs), 250 μM CuSO4 and 2.5 mM sodium ascorbate were mixed as a reaction pre-mix with 100 μM of alkyne-label: TAMRA-alkyne (Vector Labs), biotin-PEG4-alkyne (Sigma), β-GlcNAc- alkyne (Sigma). Purified protein containing pAzF was added at ∼ 1 mg/mL concentration. CuAAC reactions were rocked at RT for 1 hour.

Following either labeling procedure, labeled proteins were subsequently concentrated in 10 kDa MWCO concentrator tubes and washed with more 50 mM HEPES, 300 mM NaCl, 5% glycerol until the flowthrough ran clear, to indicate the removal of excess label. Protein aliquots were then snap frozen in liquid N2 and stored at -80C.

### Electroporation of Intein Donor proteins into mammalian cells

Several T-75 or T-175 flasks of cells were grown to >80% confluency. Cells were washed with PBS and then trypsinized and collected into a 15 mL conical tube. Cells were spun down at 100 x g for 3 minutes and media was discarded. Cells were then resuspended in 10 mL PBS, pelleted, and the supernatant discarded. This wash step was repeated once more, for a total of 2 PBS washes. Following the final wash, the cell pellet was resuspended in a low volume of “Buffer R” (ThermoFisher) or a similar-performing electroporation buffer of 250 mM sucrose, 1 mM MgCl2●6H2O in PBS^33^, such that cells were at a concentration of 4-7x10^7^/mL. Subsequently, cell:protein solutions for electroporation were made at a protein concentration of 50 µM in the solution, unless indicated otherwise. Cells were electroporated using the Neon Electroporation system (ThermoFisher). Cells were electroporated at 1400 V, 20 ms, and 2 pulses. Following electroporation, cells were transferred to tubes containing pre-warmed PBS. Cells were washed twice in PBS by centrifugation at 300 x g for 1 minute. Following washing, cells were resuspended in antibiotic-free media and added to dishes containing pre-warmed antibiotic-free media. Cells were incubated for 4 hours and then harvested by washing with PBS and then scraping in RIPA supplemented with protease inhibitors, unless otherwise specified.

### Western blotting

Cells were generally harvested by first washing with PBS and then scraping in RIPA buffer supplemented with protease inhibitors. If nuclear proteins were to be investigated, lysates were sonicated for 8 seconds (4s on, 4s off x2) at 25% power. Cell lysates were cleared by spinning at 21,000 rcf, 4 °C for 10 min. Lysate protein concentrations were normalized by BCA Assay (ThermoFisher). Lysates (typically 5-15 µg total protein per lane) were loaded onto pre-cast 4-15% SDS-PAGE gels or hand-poured 7.5% or 12% SDS-PAGE gels. Gels were run at 150V for 70 minutes and transferred onto 0.45 µM nitrocellulose membranes using a TransBlotTurbo semi-dry transfer system (BioRad). Membranes were blocked for 1 hour in 5% milk in TBST at room temp and then incubated in primary antibody in 3%BSA in TBST overnight at 4 °C. Membranes were then washed with TBST and incubated in 5% milk in TBST with secondary antibody for 1 hour at room temp. Western blots were imaged using ECL Prime reagents (Cytiva) on a GelDoc imager (BioRad).

Antibodies used: HA-HRP conjugate (#14031, CST), FLAG (#14793, CST), calnexin (533352, Novus), Myc epitope (#2276, CST), TAMRA (MA1041, Invitrogen), biotin-HRP conjugate (#7075, CST), β-actin (A3854, Sigma), 6xHis (#12698, CST), COX IV (#4850, CST), V5 (#13202, CST), c/n-Myc (#13987, CST), Chk1 (#2360, CST), and Chk1 (#32531, Abcam).

For quantification of Western blots, blots were analyzed with ImageLab (Biorad) software. Relevant bands were selected and volumes calculated using densitometry. Specifically, for edited c-Myc quantification, the edited c-Myc species band volume was normalized to the endogenous c-Myc signal from the same membrane. Half-life for edited c-Myc protein was calculated using 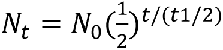

### Confocal microscopy

Cells were plated onto glass coverslips previously treated with 3.5 ug/cm^2^ CellTak (Corning). For imaging following electroporation, cells were allowed to recover and adhere to coverslips for 4 hours (or indicated time in experiment). Cells were fixed in 4% formaldehyde (ThermoFisher) for 15 minutes at room temperature, then washed 3 times with PBS. For Concanavalin A staining, cells were permeabilized in PBST (PBS with 0.25% Triton X-100) for 30 minutes at room temperature with rocking and then treated with 75 ug/mL ConA-AlexaFluor-633 (ThermoFisher) diluted in PBST for 30 minutes at room temperature. For actin cytoskeleton staining, CellMask ActinTracker Deep Red (ThermoFisher) was resuspended in DMSO to 1000x concentration, and then diluted to 1x in PBS. Cells were incubated for 15 minutes at room temperature with rocking. Cells were washed 3 times with PBS and then mounted in DAPI hard set mounting medium (Vector Labs). Coverslips were imaged on a Zeiss 980 confocal microscope.

For quantification, images were analyzed using FIJI. Images were masked using the same thresholds across the timecourse for TAMRA and DAPI, respectively. Areas above the set threshold were measured, and TAMRA signal was normalized to DAPI signal for each image.

### Flow cytometry

HAP1 cells, either WT or with CANXi14 tagged with Intein Acceptor, were electroporated with Intein Donor – TAMRA as described above. Cells were left to recover for 1 hour in antibiotic free IMDM, and then were trypsinized, pelleted in media, and then resuspended in PBS. Cell suspensions were treated with SytoxBlue (ThermoFisher) at a 1:1000 dilution, and then were filtered through a cell strainer. After a 15 minute incubation at room temperature, flow cytometry was carried out on a LSR II instrument (BD Bioscience), using a 532 nm laser line to measure TAMRA fluorescence and a 405 nm laser line to measure SytoxBlue fluorescence. Data was analyzed using web-based program Floreada.io.

### Streptavidin Pulldown of Biotinylated Protein

Lysates were added to previously equilibrated Streptavidin Agarose resin (Thermo). Lysates and resins were rocked overnight at 4 °C. Resin was pelleted by spinning at 5000 rcf for 1 minute at 4 °C. Flow throughs were collected and resin was subsequently washed 3 times with PBS. Bound proteins were eluted from Streptavidin agarose by addition of 6M guanidine-HCL, pH 1.5 followed by rapid pH neutralization. Released proteins were subsequently precipitated by ethanol precipitation by adding 9 volumes of chilled ethanol to 1 volume of eluted proteins. This mixture was left to incubate at -20 °C for 1 hour before spinning at 15,000 rpm in a chilled centrifuge. The resulting pellet was washed twice more with ice-cold ethanol before drying the pellet under vacuum to remove excess ethanol. Pellet was resuspended in loading dye or buffer for downstream applications.

### FLAG Immunoprecipitation

Lysates were added to previously equilibrated FLAG M2 resin (Sigma). Lysates and resins were rocked overnight at 4 °C. Resin was pelleted by spinning at 5000 x g for 1 minute at 4 °C. Flow throughs were collected, and resin was subsequently washed 3 times with TBS (50 mM Tris pH7.5, 150 mM NaCl) supplemented with 0.5% NP-40. Bound protein was eluted by incubating resin with 150 ng/µL FLAG epitope peptides overnight at 4 °C.

### Sample preparation for LC-MS/MS

Protein solutions were treated with 5 mM dithiothreitol (DTT) for 30 minutes at 60 °C in a heat bath, and subsequently alkylated with 20 mM iodoacetamide for 15 minutes at room temperature, in the dark. 50 μl of 50 mM TEABC pH 8.0 was added to samples before trypsin digestion was carried out using modified sequence-grade trypsin (Promega) with 1:50 trypsin:protein ratio at 37 °C overnight. Peptide desalting was carried out with the C- 18 StageTip method. The C-18 material was stacked onto 200 µl tips, activated with 100% ACN, and equilibrated with 0.1% aqueous formic acid. Peptide samples were resuspended in 0.1% formic acid and loaded onto the C-18 StageTip. The sample was passed through the tip two times, followed by washing with 0.1% formic acid and elution with 40% ACN in 0.1% formic acid. The eluent was concentrated to dryness under vacuum. The dried peptides were stored at -20 °C until LC-MS/MS analysis.

### LC-MS/MS analysis of proteins isolated from mammalian cells

The C18 cleaned peptides were analyzed on ThermoScientific Orbitrap Exploris 240 mass spectrometer interfaced with UltiMate 3000 HPLC and UHPLC Systems. Digested peptides were reconstituted in 0.1 % formic acid with a final concentration of 500 ng/µl and 1 µl of sample (500 ng) was loaded on the column. Peptides were separated on an analytical column (75 µm × 15 cm) at a flow rate of 300 nL/min using a gradient of 1–25 % solvent B (0.1 % formic acid in 100 % acetonitrile) for the first 100 minutes and 25–30 % for next 5 minutes, 30-70 % for 5 minutes, 70–1 % for next 5 minutes. The total run time was set to 120 min. The mass spectrometer was operated in data-dependent acquisition mode. A survey full scan MS (from m/z 400–1600) was acquired in the Orbitrap with a resolution of 6000 Normalized AGC target 300. Data were acquired in topN with 20 dependent scans. Fragmented used normalized collision energy of 37 % and was detected at a mass resolution of 1500. Dynamic exclusion was set for 8s with a 10 ppm mass window.

### LC-MS/MS data analysis

MS/MS searches were carried out using SEQUEST search algorithms against a Uniprot database for humans (supplemented with exogenous sequence for anticipated protein constructs) using Proteome Discoverer (Version 3.0, Thermofisher Scientific, Bremen, Germany). Oxidation of methionine and *N*-terminal protein acetylation were included as dynamic modifications and carbamidomethylation of cysteine was set as a static modification. MS and MS/MS mass tolerances were set to 10 ppm and 0.05 Da, respectively. Trypsin was specified as protease and a maximum of two missed cleavages were allowed. Target-decoy database searches used for calculation of false discovery rate (FDR) and for peptide identification FDR was set at 1%. Gene ontology (GO) analysis was carried out using PantherDB.

