## Supplemental Figures for "Intracellular Protein Editing to Enable Incorporation of Non-Canonical Residues into Endogenous Proteins"

### Supplemental Information

**Extended Data - Investigation and optimization in response to incompletely spliced product protein**

Upon comparing editing in HAP1 cells with CANX endogenously tagged with Intein Acceptor containing either mClover3 or mNeonGreen$2_{11}$, we noted that the spliced product ran at slightly different molecular weights. This observation prompted further investigation which revealed that the two Intein Acceptor constructs resulted in differential splicing activity, and thus, different spliced products (Ext. Data Fig. 1a). With electroporation and the addition of Intein Donor, both of the CANX-Intein-Acceptors containing mClover and mNG$2_{11}$ generated high molecular weight HA signal, but at different molecular weights: the edited CANX from the Intein Acceptor with mClover appears to run over 100 kDa, while the edited CANX derived from the mNG$2_{11}$ Intein Acceptor runs just below 100 kDa. Upon further investigation, the edited CANX from the mClover Intein Acceptor is also reactive to 6xHis. The 6xHis tag is situated on the recombinant Intein Donor, which indicates that the Ava:Npu split intein pair did not splice (for a detailed diagram of the tandem splicing pathway and relevant protein species, see Ext. Data Fig. 1b). Importantly, the spliced product from the mNG$2_{11}$ Intein Acceptor is not 6xHis reactive by Western blot, indicating that it is a different species and is the expected product resulting from both split intein pairs splicing.

To further verify that both Intein pairs were splicing, we created recombinant Intein Donors that were catalytically inactive for either Gp41-1, Ava:Npu, or both (Ext. Data Fig. 1c). To impair Gp41-$1_C$, essential residues Asn37 and Ser+1 were both mutated to Ala, as previously reported[1,2]. To impair $Ava_N$, the catalytic Cys1 residue was mutated to Ala. To eliminate splicing activity from both inteins, we generated an Intein Donor with both C1A (Ava:Npu impaired) and N37A/S+1A mutations (Gp41-1 impaired). This set of catalytically impaired Intein Donors were electroporated into HAP1 cells with endogenously tagged Calnexin-Int-Acceptor containing either mClover3 or mNG$2_{11}$ (Ext. Data Fig. 1d). As anticipated, the splicing results from the WT Intein Donor and the Ava:Npu impaired mutant in the mClover3-Intein Acceptor cell lines look identical, with both edited Calnexins running at the same molecular weight and both His reactive, implying that the WT Intein Donor is also not splicing with Ava:Npu. Contrastingly, with the mNG$2_{11}$ cell line there is a distinct difference between the WT ID and all impaired IDs, supporting full splicing. This experiment again confirmed that the Intein Acceptor with mNG$2_{11}$ was splicing fully, while the mClover Intein Acceptor is splicing incompletely and appears to only undergo splicing with Gp41-1.

This differential activity demonstrates that the sequence within the Intein Acceptor does have implications for splicing, and the mClover3-containing Intein Acceptor is incapable of splicing via both split intein pairs in the context of CANX. We have demonstrated that Intein Acceptors bearing the same split intein domains but different constructs separating the split inteins can have different splicing activities, and we emphasize the importance of product validation using these splicing approaches.

We reasoned that the apparent preference for the smaller mNG$2_{11}$-containing Intein Acceptor construct may be due to the large steric bulk of a mClover3 near the splicing site, and that the more rigid β-strand nature of $mNG2_{11}$ may keep the intein domains better separated with sufficient space to carry out splicing. In addition, the ~2 kDa $mNG2_{11}$ fragment has the benefit of being significantly smaller in size than mClover3, at 28 kDa, enabling an overall smaller Intein Acceptor construct. From this point forward, we proceeded with the Intein Acceptor construct containing $mNG2_{11}$ in order to ensure complete splicing in our editing approach.

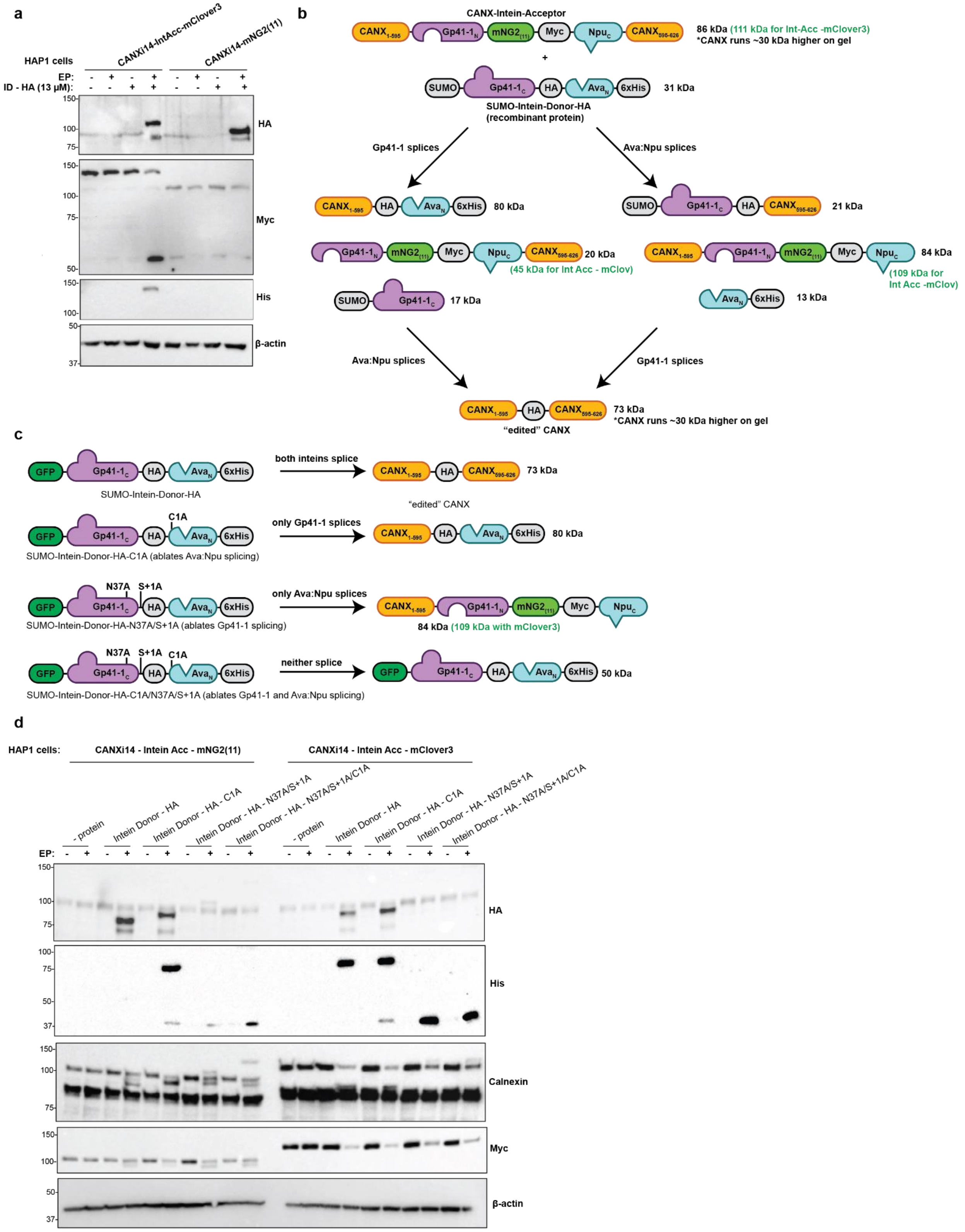

**Extended Data Figure 1 - Investigation into incomplete splicing in the dual split intein configuration. a,** Electroporation of SUMO-Intein-Donor bearing HA into HAP1 cells where intron 14 of Calnexin was tagged with Intein Acceptors, containing either mClover 3 or $mNG2_{11}$. **b,** Cartoon showing the various outcomes from tandem splicing, as the tandem reaction proceeds through one split intein pair and then the other. Approximate molecular weights of protein species are shown, with molecular weights corresponding to the CANXi14 -Intein Acceptor with mClover3 shown in paratheses in green. **c,** Cartoon of the set of catalytically inactive recombinant Intein Donors, where critical residues where mutated to alanine to disrupt splicing activity. At right, the major expected protein species, as a result of splicing, is shown. **d,** Electroporation of catalytic inactive Intein Donors from (c) into HAP1 cell lines with endogenous CANX tagged with Intein Acceptor, containing either $mNG2_{11}$ or mClover3. Not all expected protein species are discernible by Western blot.

### Supplemental Figures

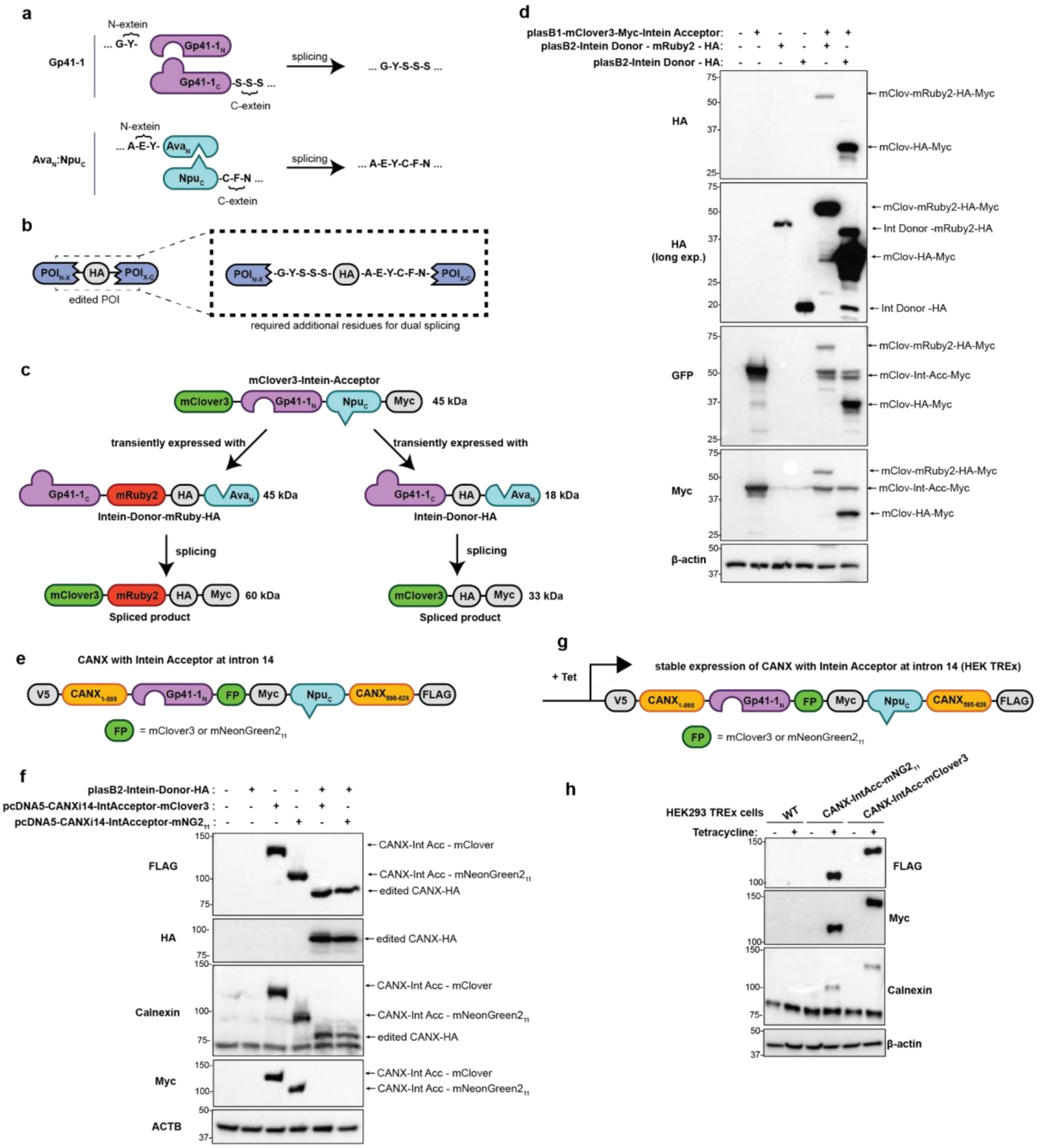

**Supplemental Figure 1 – Configuration of split intein pairs Gp41-1 and Ava:Npu for protein editing.** **a,** Gp41-1 and Ava:Npu possess different extein sequences that remain in the final spliced product. **b,** The additional residues left as a "scar" by our protein editing approach, this could be minimized in some of our model proteins. "HA" refers to an epitope tag but in this diagram, can also represent any cargo (single amino acid to larger protein sequence) to be edited into a protein of interest. **c,** A schematic for the concatenation of fluorescent proteins and epitope tags by tandem split intein mediated protein trans-splicing. **d,** Western blots resulting from the transient expression of constructs shown in (B) in HEK293T cells. **e,** Design of calnexin – Intein Acceptor constructs that contain a Myc epitope tag and either an $mNG2_{11}$ or mClover3. **f,** Transient expression of two calnexin – Intein Acceptor sequences containing either mClover3 or $mNG2_{11}$ in combination with transient expression of an Intein donor bearing HA, resulting in Calnexin edited to contain an HA epitope at intron 14. **g,** Schematic of calnexin – Intein Acceptor construct for stable HEK293 TREx cells**. h,** Generation of stable HEK293 TREx cells that express a FLAG-tagged calnexin – Intein Acceptor construct upon induction with Tetracycline. The Intein Acceptor either contains mClover3 or a smaller $mNG2_{11}$.

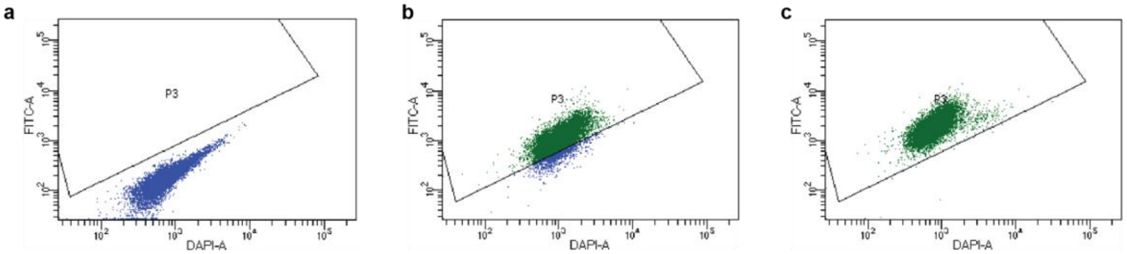

**Supplemental Figure 2 – Generation of endogenously tagged proteins using an intron tagging Cas9 approach. a,** Flow cytometry for parent HAP1 cells, showing low levels of GFP signal from cells. **b,** Flow cytometry for HAP1 cells with endogenous Calnexin tagged at intron 14 with Intein Acceptor containing $mNG2_{11}$ with gate demonstrating increased levels of GFP signal. **c,** Flow cytometry for HAP1 cells with endogenous Calnexin tagged at intron 14 with Intein Acceptor containing mClover3 with gate demonstrating increased levels of GFP signal.

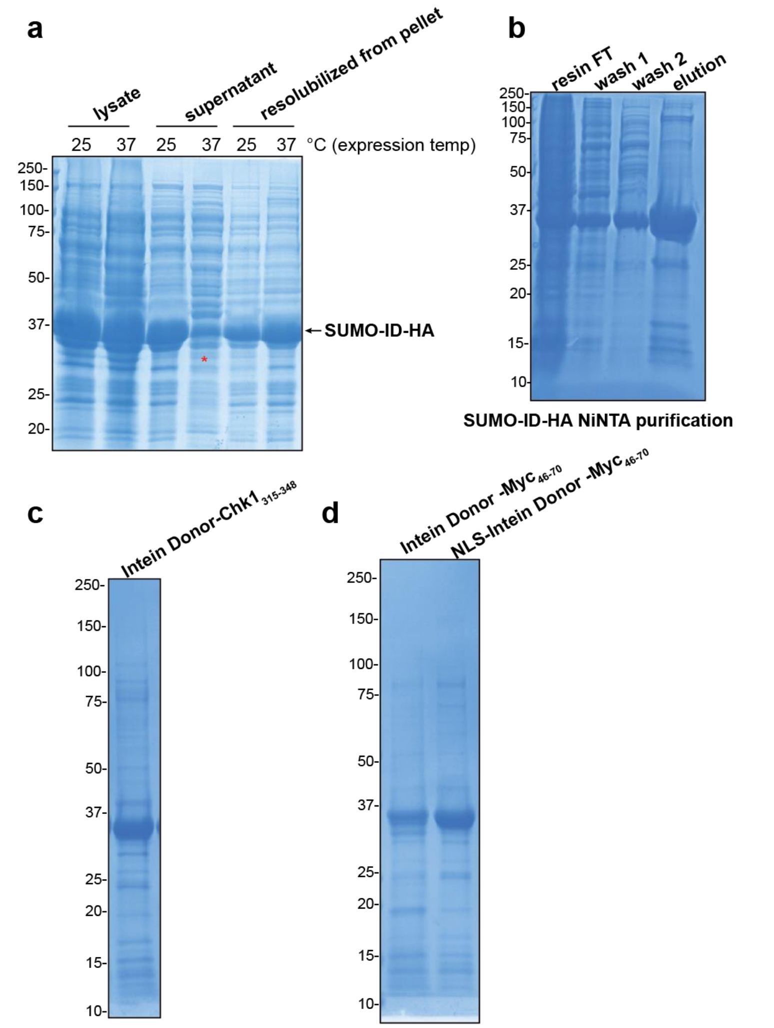

**Supplemental Figure 3- Expression and Purification of Recombinant Intein Donor.**
**a,** Intein Donor bearing HA epitope tag expression is sensitive to expression temperature. At 37°C, the protein mostly accumulates in the pellet (red asterisk), while at 25°C the protein is partitioned between both the supernatant and pellet. **b,** Intein Donor bearing HA epitope tag purification by NiNTA resin, following resolubilization from the pellet. **c,** Purified Intein Donor bearing Chk1$_{315\text{-}348}$. **d,** Purified Intein Donors bearing c-Myc$_{46\text{-}70}$, either with or without an NLS sequence.

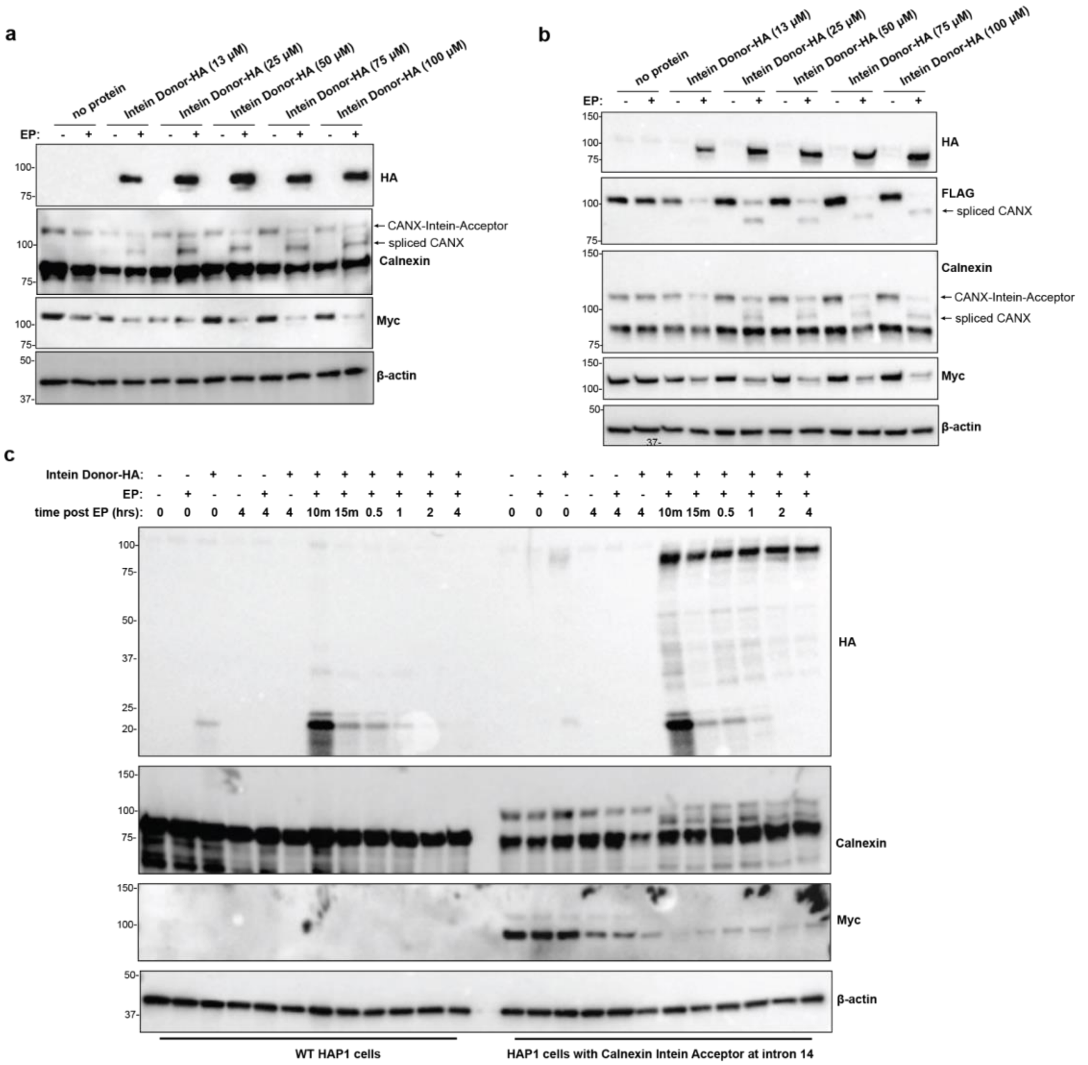

**Supplemental Figure 4 - Preliminary data for protein editing in Calnexin. a,** Electroporation of a variety of concentrations of recombinant Intein Donor into HAP1 cells with endogenous Calnexin tagged with Intein Acceptor $mNG2_{11}$ at intron 14. The concentration of Intein Donor refers to the amount present in the electroporation solution, prior to electroporation. **b,** Electroporation of a variety of concentrations of recombinant Intein Donor into HEK293 cells stably expressing FLAG-tagged Calnexin with Intein Acceptor $mNG2_{11}$ at intron 14. The concentration of Intein Donor refers to the amount present in the electroporation solution, prior to electroporation. **c,** Electroporation of Intein Donor – HA into WT HAP1 cells or HAP1 cells with Calnexin – Intein Acceptor, with samples harvested at timepoints following electroporation and visualized by Western blot.

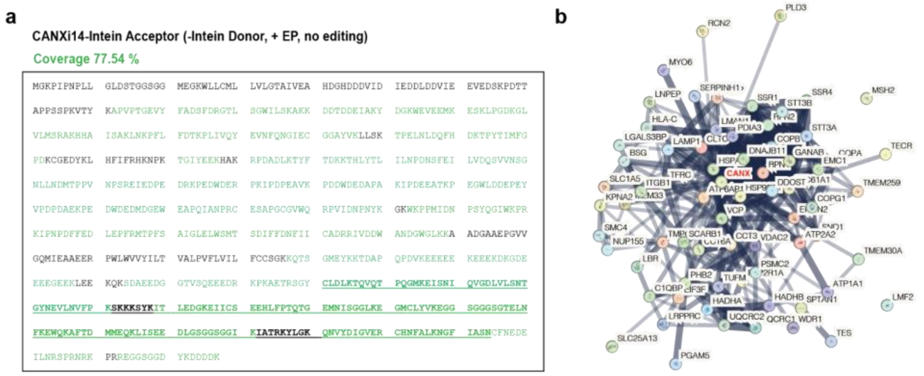

**Supplemental Figure 5 - Mass spectrometry of edited Calnexin from stable HEK293 TREx cells. a,** Sequence coverage of FLAG-tagged Calnexin -Intein Acceptor (with no editing) from HEK293 TREx cells, as identified by mass spectrometry following FLAG pulldown and trypsin digestion. The underlined region is the Intein Acceptor sequence. Residues in green correspond to peptides detected by MS. **b,** A network of the 70 proteins shared between our dataset and the Biogrid database, shown with known interactions between proteins.

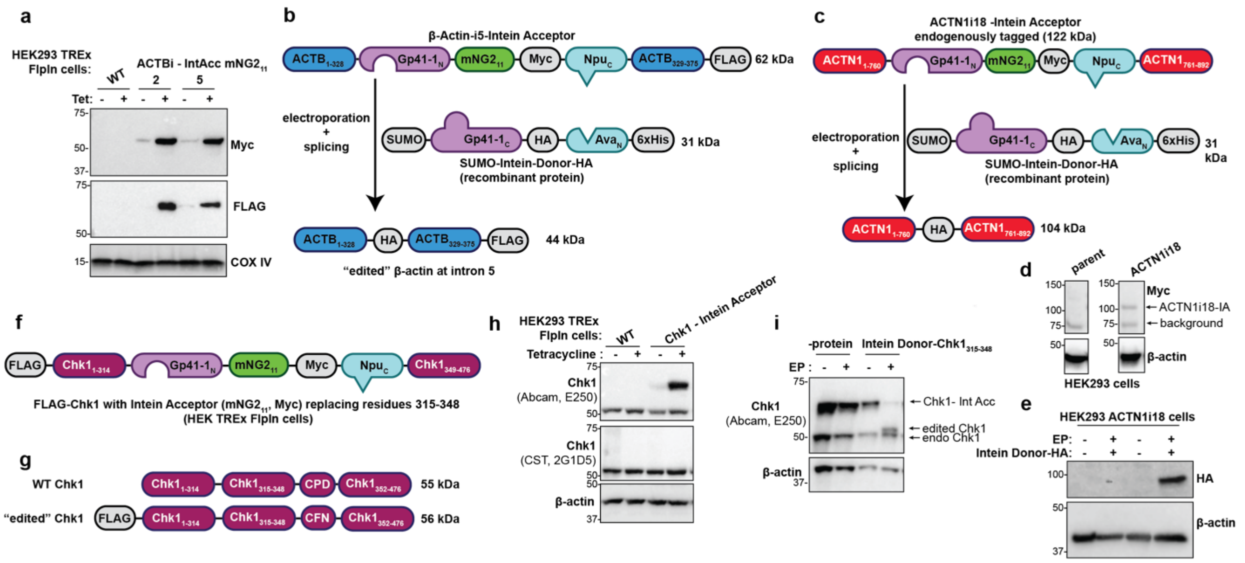

**Supplemental Figure 6 - Protein editing in β-actin, α-actinin-1, and Chk1. a,** Generation of HEK293 TREx cell lines stably expressing β-actin with Intein Acceptor. **b,** Schematic of β-actin editing at intron 5, in HEK293 TREx cells stably expressing β-actin with Intein Acceptor at intron 5. **c,** Schematic of endogenous α-actinin-1 editing at intron 18. **d,** Generation of a HEK293 cell line where α-actinin-1 has been endogenously tagged with Intein Acceptor at intron 18, in comparison to the parent cell line. **e,** Editing α-actinin-1 to include an HA epitope tag at the site corresponding to intron 18, using the endogenously tagged cell line from (D). **f,** Schematic of Chk1- Intein Acceptor construct, where Intein Acceptor replaces residues 315 – 348. **g,** Comparison of edited Chk1 to WT Chk1, with only 2 point mutations required to accommodate extein residues**. h,** Generation of stable HEK293 TREx lines that express Chk1-Intein Acceptor, containing $mNG2_{11}$ and a Myc epitope tag, upon induction with Tetracycline**. i,** Chk1 editing in response to electroporation and Intein Donor bearing $Chk1_{315-348}$ at 1 hour post-electroporation.

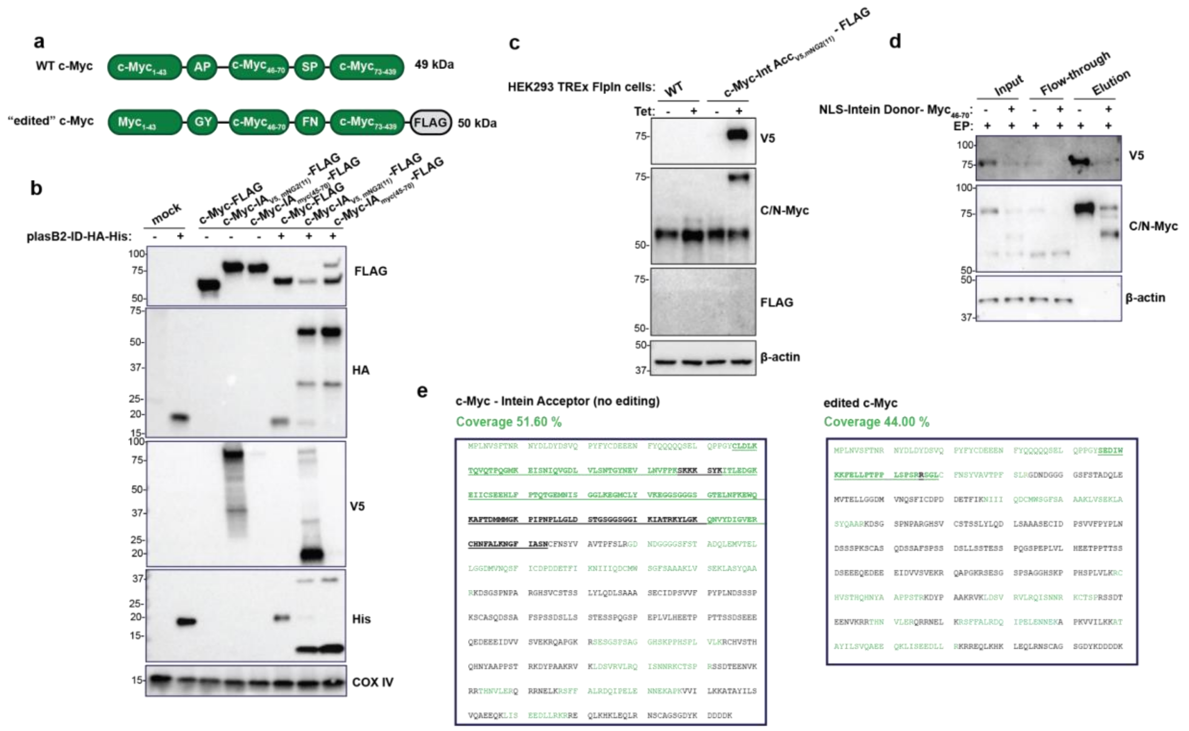

**Supplemental Figure 7 - Applying protein editing method to c-Myc.** **a,** Comparison of WT and edited c-Myc, showing that edited c-Myc has point mutations to accommodate extein residues. **b,** Transient expression of several c-Myc-Intein Acceptor constructs and Intein Donor bearing HA results in edited c-Myc in HEK293T cells. **c,** Generation of stable HEK293 TREx lines that express c-Myc-Intein Acceptor, containing $mNG2_{11}$ and a V5 epitope tag, upon induction with Tetracycline. **d,** Pulldown of either c-Myc-Intein Acceptor or edited c-Myc using FLAG resin demonstrates complete, dual-spliced c-Myc containing a C-terminal FLAG tag, despite the lack of FLAG signal on Western blots. **e,** Sequence coverage, as determined by mass spectrometry, of the c-Myc -Intein Acceptor or edited c-Myc proteins isolated by the FLAG pulldown in (D). The underlined region is the Intein Acceptor sequence (left) or the region that was edited in (right). Residues in green correspond to peptides detected by MS.

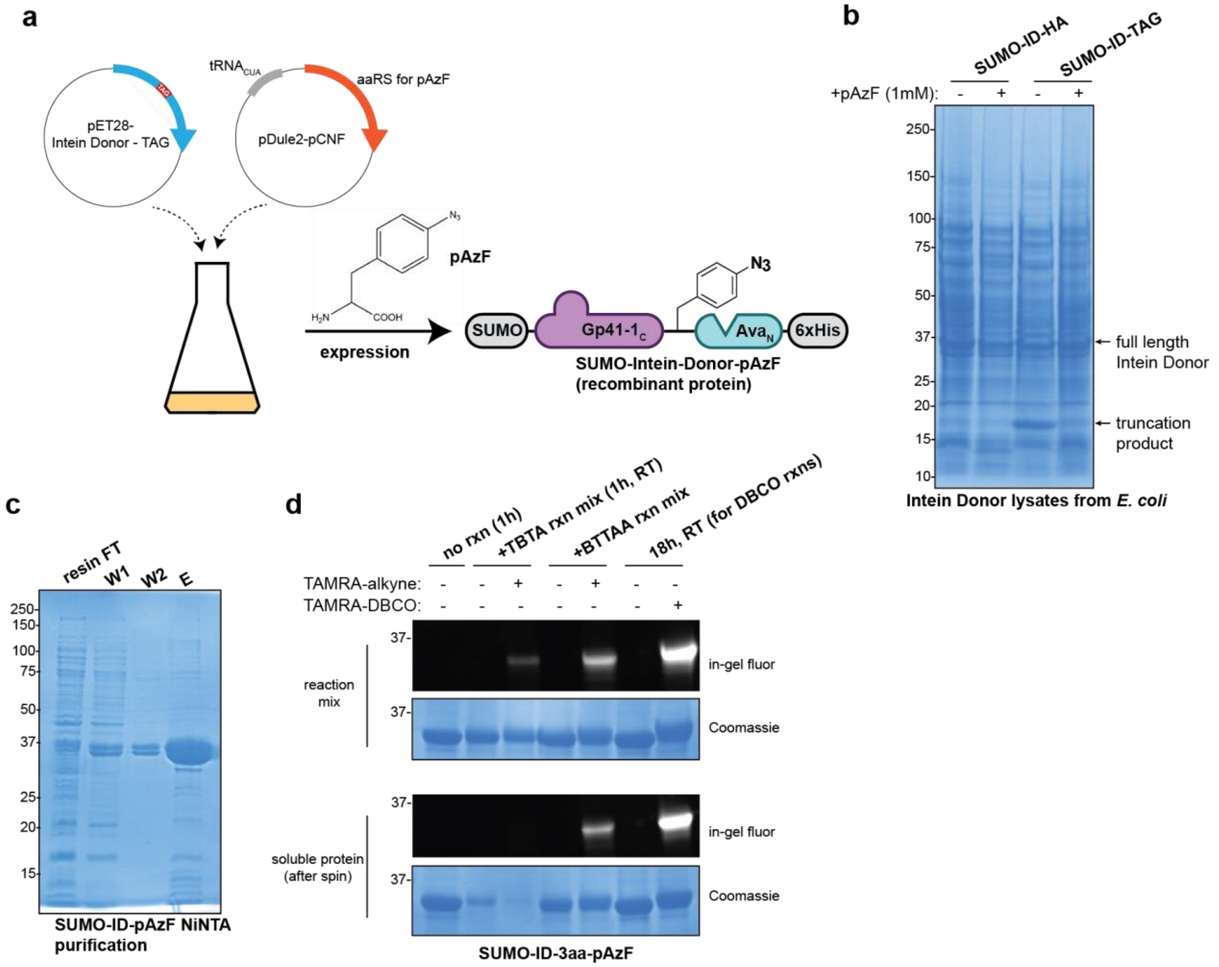

**Supplemental Figure 8 – Genetic code expansion in *E. coli* enables recombinant Intein Donors containing useful and diverse functional groups. a,** Genetic code expansion (GCE) enables the site specific encoding of ncAAs into proteins, shown here in *E. coli*. In this way, we can produce Intein Donor containing a pAzF ncAA. **b,** Lysates from a GCE trial expression show production of full-length Intein Donor with pAzF only in the presence of 1 mM pAzF (without pAzF, truncation product is formed). **c,** NiNTA purification for recombinant Intein Donor bearing pAzF. **d,** Conjugating labels onto purified Intein Donor – pAzF with CuAAC (using TBTA or BTTAA as ligand) or Cu-free SPAAC click chemistry shows varying degrees of protein solubility following labelling.

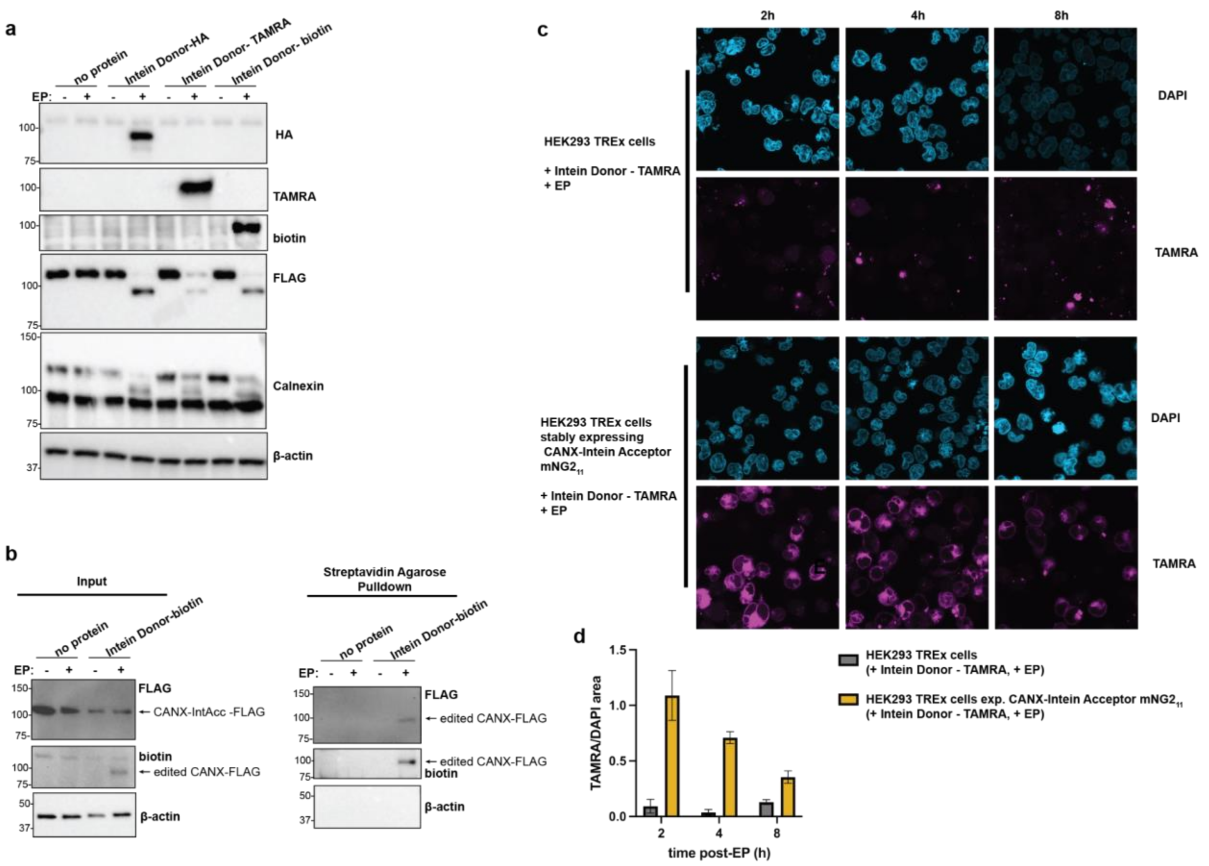

**Supplemental Figure 9** - **Additional data for editing pAzF and labels into Calnexin in HEK293 stable cell lines. a,** Editing HA tag, TAMRA, and biotin labels into Calnexin in stable HEK293 TREx line. **b,** Editing of biotin into Calnexin in the stable HEK293 TREx cell line followed by streptavidin pulldown results in isolation of the edited, biotinylated protein. **c,** Confocal microscopy timecourse following electroporation of Intein Donor -TAMRA in either WT HEK293 TREx cells or HEK293 TREx cells with Calnexini14-Intein Acceptor-$mNG2_{11}$. **d,** Quantification of the area of TAMRA signal normalized to DAPI signal for the timecourse in (C) (n=3 or more for each condition)

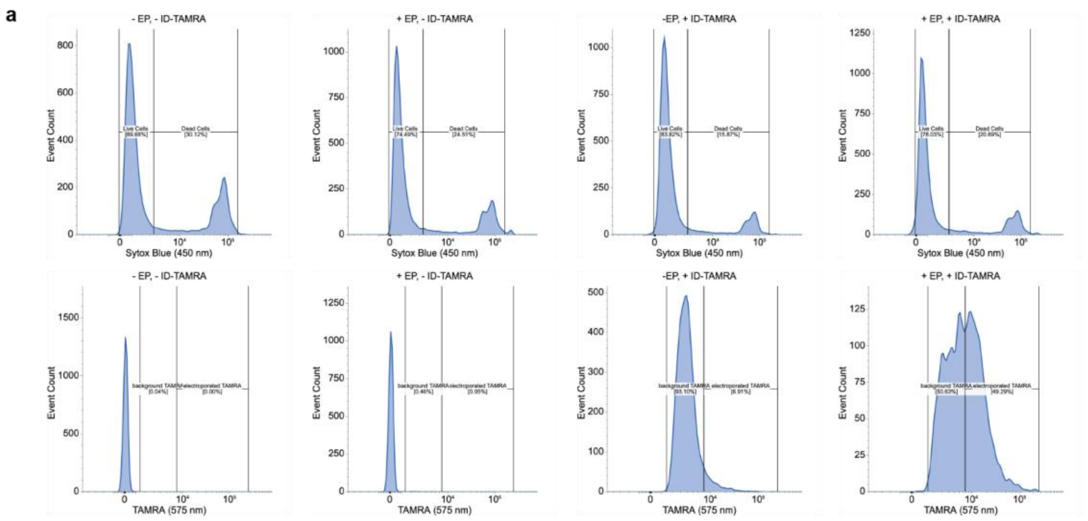

**Supplemental Figure 10 – Flow cytometry of cells post-electroporation with Intein Donor – TAMRA supports robust delivery. a,** Flow cytometry following electroporation of Intein Donor bearing TAMRA as cargo into HAP1 cells with endogenous Calnexin tagged with Intein Acceptor – mNG2$_{11}$ at intron 14. The upper row shows various conditions stained with Sytox Blue (stains dead cells) as a histogram. The lower row shows the TAMRA signal from various conditions as a histogram.

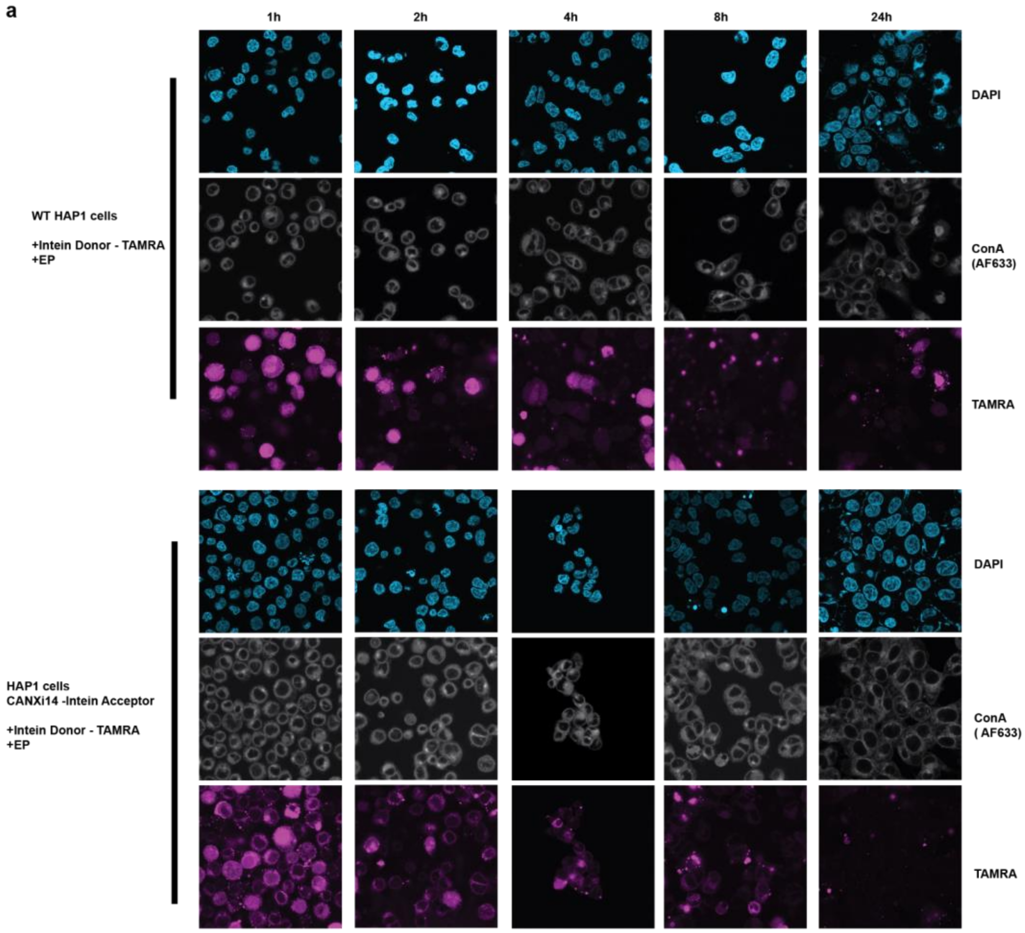

**Supplemental Figure 11 - Imaging timecourse for Calnexin edited to include TAMRA, with ConA as an ER marker. a,** Confocal microscopy timecourse following electroporation of Intein Donor -TAMRA into either WT HAP1 cells or HAP1 cells with CANXi14 endogenously tagged with Intein Acceptor-mNG2(11), with Concavalin A – AlexaFluor 633 for ER marker.

### Supplemental Tables

Table S1 – Calnexin Co-IP Protein List
Table S2 – Acceptor Myc Co-IP Protein List
Table S3 – Edited Myc Co-IP Protein

### Protein Sequences

#### Preliminary Transient Expression Constructs

Key:
mClover3
mRuby
Gp41-1$_{C}$
Ava$_{N}$
Gp41-1$_{N}$
Npu$_{C}$
**Myc (bold)**
HA

**plasB1-mClover3-Myc-Intein Acceptor**

MVSKGEELFTGVVPILVELDGDVNGHKFSVRGEGEGDATNGKLTLKFICTTGKLPVPWPTLVTT
FGYGVACFSRYPDHMKQHDFFKSAMPEGYVQERTISFKDDGTYKTRAEVKFEGDTLVNRIELKG
IDFKEDGNILGHKLEYNFNSHYVYITADKQKNCIKANFKIRHNVEDGSVQLADHYQQNTPIGDG
PVLLPDNHYLSHQSKLSKDPNEKRDHMVLLEFVTAAGITHGMDELYKGSGGSGTRSGYCLDLKT
QVQTPQGMKEISNIQVGDLVLSNTGYNEVLNVFPKSKKKSYKITLEDGKEIICSEEHLFPTQTG
EMNISGGLKEGMCLYVKEGGSGGGSGGSGGSGGIKIATRKYLGKQNVYDIGVERCHNFALKNGF
IASNCFNGSGG**EQKLISEEDL***

**plasB2-Intein Donor - HA**

MCLKKILKIEELDERELIDIEVCGNHLFYANDILTHNSSSYPYDVPDYAAEYCLSYDTEVLTVE
YGFVPIGEIVDKGIECSVFSIDSNGIVYTQPIAQWHHRGKQEVFEYCLEDGSIIKATKDHKFMT
QDGKMLPIDEIFEQELDLLQVKGLPE*

**plasB2-Intein Donor - mRuby - HA**

MCLKKILKIEELDERELIDIEVCGNHLFYANDILTHNSSSDVGSSVSKGEELIKENMRMKVVME
GSVNGHQFKCTGEGEGRPYEGVQTMRIKVIEGGPLPFAFDILATSFMYGSRTFIKYPADIPDFF
KQSFPEGFTWERVTRYEDGGVVTVTQDTSLEDGELVYNVKVRGVNFPSNGPVMQKKTKGWEPNT
EMMYPADGGLRGYTDIALKVDGGGHLHCNFVTTYRSKKTVGNIKMPGVHAVDHRLERIEESDNE
TYVVQREVAVAKYSNLGGGMDELYKYPYDVPDYAGGSAEYCLSYDTEVLTVEYGFVPIGEIVDK
GIECSVFSIDSNGIVYTQPIAQWHHRGKQEVFEYCLEDGSIIKATKDHKFMTQDGKMLPIDEIF
EQELDLLQVKGLPE*

#### Relevant Recombinant Protein Sequences

Key:
SUMO

Peptide/cargo to be spliced into target protein- Intein Acceptor
Gp41-1C
AvaN
Various catalytic mutations
NLS

**SUMO-Gp41-1C-HA-AvaN-6xHis:**

MSDSEVNQEAKPEVKPEVKPETHINLKVSDGSSEIFFKIKKTTPLRRLMEAFAKRQGKEMDSLR
FLYDGIRIQADQTPEDLDMEDNDIIEAHREQIGGGGSGGSGGLVPRGSGGSGGMCLKKILKIEE
LDERELIDIEVCGNHLFYANDILTHNSSSYPYDVPDYAAEYCLSYDTEVLTVEYGFVPIGEIVD
KGIECSVFSIDSNGIVYTQPIAQWHHRGKQEVFEYCLEDGSIIKATKDHKFMTQDGKMLPIDEI
FEQELDLLQVKGLPELEHHHHHH*

**SUMO-Gp41-1C-pAzF-AvaN-6xHis:**

MSDSEVNQEAKPEVKPEVKPETHINLKVSDGSSEIFFKIKKTTPLRRLMEAFAKRQGKEMDSLR
FLYDGIRIQADQTPEDLDMEDNDIIEAHREQIGGGGSGGSGGLVPRGSGGSGGMCLKKILKIEE
LDERELIDIEVCGNHLFYANDILTHNSSS(pAzF)AEYCLSYDTEVLTVEYGFVPIGEIVDKGI
ECSVFSIDSNGIVYTQPIAQWHHRGKQEVFEYCLEDGSIIKATKDHKFMTQDGKMLPIDEIFEQ
ELDLLQVKGLPELEHHHHHH*

**GFP-Gp41-1C-HA-AvaN-6xHis (C1A):**

MVSKGEELFTGVVPILVELDGDVNGHKFSVRGEGEGDATNGKLTLKFICTTGKLPVPWPTLVTT
LTYGVQCFSRYPDHMKRHDFFKSAMPEGYVQERTISFKDDGTYKTRAEVKFEGDTLVNRIELKG
IDFKEDGNILGHKLEYNFNSHNVYITADKQKNGIKANFKIRHNVEDGSVQLADHYQQNTPIGDG
PVLLPDNHYLSTQSVLSKDPNEKRDHMVLLEFVTAAGITHGMDELYKGSMCLKKILKIEELDER
ELIDIEVCGNHLFYANDILTHNSSSYPYDVPDYAAEYALSYDTEVLTVEYGFVPIGEIVDKGIE
CSVFSIDSNGIVYTQPIAQWHHRGKQEVFEYCLEDGSIIKATKDHKFMTQDGKMLPIDEIFEQE
LDLLQVKGLPELEHHHHHH*

**GFP-Gp41-1C-HA-AvaN-6xHis (N37A/S+1A):**

MVSKGEELFTGVVPILVELDGDVNGHKFSVRGEGEGDATNGKLTLKFICTTGKLPVPWPTLVTT
LTYGVQCFSRYPDHMKRHDFFKSAMPEGYVQERTISFKDDGTYKTRAEVKFEGDTLVNRIELKG
IDFKEDGNILGHKLEYNFNSHNVYITADKQKNGIKANFKIRHNVEDGSVQLADHYQQNTPIGDG
PVLLPDNHYLSTQSVLSKDPNEKRDHMVLLEFVTAAGITHGMDELYKGSMCLKKILKIEELDER
ELIDIEVCGNHLFYANDILTHAASSYPYDVPDYAAEYCLSYDTEVLTVEYGFVPIGEIVDKGIE
CSVFSIDSNGIVYTQPIAQWHHRGKQEVFEYCLEDGSIIKATKDHKFMTQDGKMLPIDEIFEQE
LDLLQVKGLPELEHHHHHH*

**GFP-Gp41-1C-HA-AvaN-6xHis (C1A/N37A/S+1A):**

MVSKGEELFTGVVPILVELDGDVNGHKFSVRGEGEGDATNGKLTLKFICTTGKLPVPWPTLVTT
LTYGVQCFSRYPDHMKRHDFFKSAMPEGYVQERTISFKDDGTYKTRAEVKFEGDTLVNRIELKG
IDFKEDGNILGHKLEYNFNSHNVYITADKQKNGIKANFKIRHNVEDGSVQLADHYQQNTPIGDG
PVLLPDNHYLSTQSVLSKDPNEKRDHMVLLEFVTAAGITHGMDELYKGSMCLKKILKIEELDER
ELIDIEVCGNHLFYANDILTHAASSYPYDVPDYAAEYALSYDTEVLTVEYGFVPIGEIVDKGIE
CSVFSIDSNGIVYTQPIAQWHHRGKQEVFEYCLEDGSIIKATKDHKFMTQDGKMLPIDEIFEQE
LDLLQVKGLPELEHHHHHH*

**SUMO- Gp41-1$_C$-Chk1$_{315-348}$-Ava$_N$-6xHis:**

MSDSEVNQEAKPEVKPEVKPETHINLKVSDGSSEIFFKIKKTTPLRRLMEAFAKRQGKEMDSLR
FLYDGIRIQADQTPEDLDMEDNDIIEAHREQIGGGGSGGSGGLVPRGSGGSGGMCLKKILKIEE
LDERELIDIEVCGNHLFYANDILTHNSSSQPEPRTGLSLWDTSPSYIDKLVQGISFSQPTCLSY
DTEVLTVEYGFVPIGEIVDKGIECSVFSIDSNGIVYTQPIAQWHHRGKQEVFEYCLEDGSIIKA
TKDHKFMTQDGKMLPIDEIFEQELDLLQVKGLPELEHHHHHH*

**SUMO-NLS-Gp41-1$_C$-Myc$_{40-75}$-Ava$_N$-6xHis:**

MSDSEVNQEAKPEVKPEVKPETHINLKVSDGSSEIFFKIKKTTPLRRLMEAFAKRQGKEMDSLR
FLYDGIRIQADQTPEDLDMEDNDIIEAHREQIGGGGSGGSGGPKKKRKVGSGGSGGMCLKKILK
IEELDERELIDIEVCGNHLFYANDILTHNSEDIWKKFELLPTPPLSPSRRSGLCLSYDTEVLTV
EYGFVPIGEIVDKGIECSVFSIDSNGIVYTQPIAQWHHRGKQEVFEYCLEDGSIIKATKDHKFM
TQDGKMLPIDEIFEQELDLLQVKGLPELEHHHHHH*

---

### Protein Sequences from stable HEK293 TREx FlpIn mammalian cell lines (in a pcDNA5 vector to generate stable cell line)

Key:
V5
FLAG
Gp41-1$_N$
Npu$_C$
Myc epitope tag
Fluorescent protein (mNG2$_{11}$ or mClover)
Residue mutated from WT for extein
**Bold = residues inserted for extein**

**V5-CANXi14- Intein Acceptor (mNG2$_{11}$, Myc)-FLAG**

MGKPIPNPLLGLDSTGGSGGMEGKWLLCMLLVLGTAIVEAHDGHDDDVIDIEDDLDDVIEEVED
SKPDTTAPPSSPKVTYKAPVPTGEVYFADSFDRGTLSGWILSKAKKDDTDDEIAKYDGKWEVEE
MKESKLPGDKGLVLMSRAKHHAISAKLNKPFLFDTKPLIVQYEVNFQNGIECGGAYVKLLSKTP
ELNLDQFHDKTPYTIMFGPDKCGEDYKLHFIFRHKNPKTGIYEEKHAKRPDADLKTYFTDKKTH
LYTLILNPDNSFEILVDQSVVNSGNLLNDMTPPVNPSREIEDPEDRKPEDWDERPKIPDPEAVK
PDDWDEDAPAKIPDEEATKPEGWLDDEPEYVPDPDAEKPEDWDEDMDGEWEAPQIANPRCESAP
GCGVWQRPVIDNPNYKGKWKPPMIDNPSYQGIWKPRKIPNPDFFEDLEPFRMTPFSAIGLELWS
MTSDIFFDNFIICADRRIVDDWANDGWGLKKAADGAAEPGVVGQMIEAAEERPWLWVVYILTVA
LPVFLVILFCCSGKKQTSGMEYKKTDAPQPDVKEEEEEKEEEKDKGDEEEEGEEKLEEKQKSDA
EEDGGTVSQEEEDRKPKAE**TRSGY**CLDLKTQVQTPQGMKEISNIQVGDLVLSNTGYNEVLNVFP
KSKKKSYKITLEDGKEIICSEEHLFPTQTGEMNISGGLKEGMCLYVKEGGSGGGSGTELNFKEW
QKAFTDMMEQKLISEEDLGSGGSGGIKIATRKYLGKQNVYDIGVERCHNFALKNGFIASN**CFN**E
DEILNRSPRNRKPRREGGSGGDYKDDDDK*

**V5-CANXi14- Intein Acceptor (mClover3, Myc)-FLAG**

MGKPIPNPLLGLDSTGGSGGMEGKWLLCMLLVLGTAIVEAHDGHDDDVIDIEDDLDDVIEEVED
SKPDTTAPPSSPKVTYKAPVPTGEVYFADSFDRGTLSGWILSKAKKDDTDDEIAKYDGKWEVEE

MKESKLPGDKGLVLMSRAKHHAISAKLNKPFLFDTKPLIVQYEVNFQNGIECGGAYVKLLSKTP
ELNLDQFHDKTPYTIMFGPDKCGEDYKLHFIFRHKNPKTGIYEEKHAKRPDADLKTYFTDKKTH
LYTLILNPDNSFEILVDQSVVNSGNLLNDMTPPVNPSREIEDPEDRKPEDWDERPKIPDPEAVK
PDDWDEDAPAKIPDEEATKPEGWLDDEPEYVPDPDAEKPEDWDEDMDGEWEAPQIANPRCESAP
GCGVWQRPVIDNPNYKGKWKPPMIDNPSYQGIWKPRKIPNPDFFEDLEPFRMTPFSAIGLELWS
MTSDIFFDNFIICADRRIVDDWANDGWGLKKAADGAAEPGVVGQMIEAAEERPWLWVVYILTVA
LPVFLVILFCCSGKKQTSGMEYKKTDAPQPDVKEEEEEKEEEKDKGDEEEEGEEKLEEKQKSDA
EEDGGTVSQEEEDRKPKAE**TRSGY**CLDLKTQVQTPQGMKEISNIQVGDLVLSNTGYNEVLNVFP
KSKKKSYKITLEDGKEIICSEEHLFPTQTGEMNISGGLKEGMCLYVKEGGSGGGSGVSKGEELF
TGVVPILVELDGDVNGHKFSVRGEGEGDATNGKLTLKFICTTGKLPVPWPTLVTTFGYGVACFS
RYPDHMKQHDFFKSAMPEGYVQERTISFKDDGTYKTRAEVKFEGDTLVNRIELKGIDFKEDGNI
LGHKLEYNFNSHYVYITADKQKNCIKANFKIRHNVEDGSVQLADHYQQNTPIGDGPVLLPDNHY
LSHQSKLSKDPNEKRDHMVLLEFVTAAGITHGMDELYKEQKLISEEDLGSGGSGGIKIATRKYL
GKQNVYDIGVERCHNFALKNGFIASN**CFN**EDEILNRSPRNRKPRREGGSGGDYKDDDDK*

**ACTBi2-Intein Acceptor (mNG2$_{11}$, Myc) – FLAG**

MDDDIAALVVDNGSGMCKAGFAGDDAPRAVFPSIVGRPRHQ**GY**CLDLKTQVQTPQGMKEISNIQ
VGDLVLSNTGYNEVLNVFPKSKKKSYKITLEDGKEIICSEEHLFPTQTGEMNISGGLKEGMCLY
VKEGGSGGGSGTELNFKEWQKAFTDMMEQKLISEEDLGSGGSGGIKIATRKYLGKQNVYDIGVE
RCHNFALKNGFIASN**CFN**GVMVGMGQKDSYVGDEAQSKRGILTLKYPIEHGIVTNWDDMEKIWH
HTFYNELRVAPEEHPVLLTEAPLNPKANREKMTQIMFETFNTPAMYVAIQAVLSLYASGRTTGI
VMDSGDGVTHTVPIYEGYALPHAILRLDLAGRDLTDYLMKILTERGYSFTTTAEREIVRDIKEK
LCYVALDFEQEMATAASSSSLEKSYELPDGQVITIGNERFRCPEALFQPSFLGMESCGIHETTF
NSIMKCDVDIRKDLYANTVLSGGTTMYPGIADRMQKEITALAPSTMKIKIIAPPERKYSVWIGG
SILASLSTFQQMWISKQEYDESGPSIVHRKCFGGSGGDYKDDDDK*

**ACTBi5-Intein Acceptor (mNG2$_{11}$, Myc) – FLAG**

MDDDIAALVVDNGSGMCKAGFAGDDAPRAVFPSIVGRPRHQGVMVGMGQKDSYVGDEAQSKRGI
LTLKYPIEHGIVTNWDDMEKIWHHTFYNELRVAPEEHPVLLTEAPLNPKANREKMTQIMFETFN
TPAMYVAIQAVLSLYASGRTTGIVMDSGDGVTHTVPIYEGYALPHAILRLDLAGRDLTDYLMKI
LTERGYSFTTTAEREIVRDIKEKLCYVALDFEQEMATAASSSSLEKSYELPDGQVITIGNERFR
CPEALFQPSFLGMESCGIHETTFNSIMKCDVDIRKDLYANTVLSGGTTMYPGIADRMQKEITAL
APSTMKIK**GY**CLDLKTQVQTPQGMKEISNIQVGDLVLSNTGYNEVLNVFPKSKKKSYKITLEDG
KEIICSEEHLFPTQTGEMNISGGLKEGMCLYVKEGGSGGGSGTELNFKEWQKAFTDMMEQKLIS
EEDLGSGGSGGIKIATRKYLGKQNVYDIGVERCHNFALKNGFIASN**CFN**IIAPPERKYSVWIGG
SILASLSTFQQMWISKQEYDESGPSIVHRKCFGGSGGDYKDDDDK*

**c-Myc – Intein Acceptor (mNG2$_{11}$, V5) – FLAG (A44G,P45Y, S71F, P72N)**

MPLNVSFTNRNYDLDYDSVQPYFYCDEEENFYQQQQQSELQPPGYCLDLKTQVQTPQGMKEISN
IQVGDLVLSNTGYNEVLNVFPKSKKKSYKITLEDGKEIICSEEHLFPTQTGEMNISGGLKEGMC
LYVKEGGSGGGSGTELNFKEWQKAFTDMMGKPIPNPLLGLDSTGSGGSGGIKIATRKYLGKQN
VYDIGVERCHNFALKNGFIASNCFNSYVAVTPFSLRGDNDGGGGSFSTADQLEMVTELLGGDMV
NQSFICDPDDETFIKNIIIQDCMWSGFSAAAKLVSEKLASYQAARKDSGSPNPARGHSVCSTSS
LYLQDLSAAASECIDPSVVFPYPLNDSSSPKSCASQDSSAFSPSSDSLLSSTESSPQGSPEPLV
LHEETPPTTSSDSEEEQEDEEEIDVVSVEKRQAPGKRSESGSPSAGGHSKPPHSPLVLKRCHVS
THQHNYAAPPSTRKDYPAAKRVKLDSVRVLRQISNNRKCTSPRSSDTEENVKRRTHNVLERQRR

NELKRSFFALRDQIPELENNEKAPKVVILKKATAYILSVQAEEQKLISEEDLLRKRREQLKHKL
EQLRNSCAGSGDYKDDDDK*

**Chk1 – Intein Acceptor (mNG2$_{11}$, Myc) – FLAG (P350F, D351N)**

MGDYKDDDDKMAVPFVEDWDLVQTLGEGAYGEVQLAVNRVTEEAVAVKIVDMKRAVDCPENIKK
EICINKMLNHENVVKFYGHRREGNIQYLFLEYCSGGELFDRIEPDIGMPEPDAQRFFHQLMAGV
VYLHGIGITHRDIKPENLLLDERDNLKISDFGLATVFRYNNRERLLNKMCGTLPYVAPELLKRR
EFHAEPVDVWSCGIVLTAMLAGELPWDQPSDSCQEYSDWKEKKTYLNPWKKIDSAPLALLHKIL
VENPSARITIPDIKKDRWYNKPLKKGAKRPRVTSGGVSESPSGFSKHIQSNLDFSPVNSASSEE
NVKYCLDLKTQVQTPQGMKEISNIQVGDLVLSNTGYNEVLNVFPKSKKKSYKITLEDGKEIICS
EEHLFPTQTGEMNISGGLKEGMCLYVKEGGSGGGSGTELNFKEWQKAFTDMMEQKLISEEDLGS
GGSGGIKIATRKYLGKQNVYDIGVERCHNFALKNGFIASNCFNHMLLNSQLLGTPGSSQNPWQR
LVKRMTRFFTKLDADKSYQCLKETCEKLGYQWKKSCMNQVTISTTDRRNNKLIFKVNLLEMDDK
ILVDFRLSKGDGLEFKRHFLKIKGKLIDIVSSQKVWLPAT*

---

**Protein Sequences from endogenously tagged proteins**

Key:
**Bold = residues inserted for extein**
Gp41-1$_N$
Npu$_C$
Myc epitope tag
Fluorescent protein (mNG2$_{11}$ or mClover)

**CANXi14- Intein Acceptor (mClover3, Myc)**

MEGKWLLCMLLVLGTAIVEAHDGHDDDVIDIEDDLDDVIEEVEDSKPDTTAPPSSPKVTYKAPV
PTGEVYFADSFDRGTLSGWILSKAKKDDTDDEIAKYDGKWEVEEMKESKLPGDKGLVLMSRAKH
HAISAKLNKPFLFDTKPLIVQYEVNFQNGIECGGAYVKLLSKTPELNLDQFHDKTPYTIMFGPD
KCGEDYKLHFIFRHKNPKTGIYEEKHAKRPDADLKTYFTDKKTHLYTLILNPDNSFEILVDQSV
VNSGNLLNDMTPPVNPSREIEDPEDRKPEDWDERPKIPDPEAVKPDDWDEDAPAKIPDEEATKP
EGWLDDEPEYVPDPDAEKPEDWDEDMDGEWEAPQIANPRCESAPGCGVWQRPVIDNPNYKGKWK
PPMIDNPSYQGIWKPRKIPNPDFFEDLEPFRMTPFSAIGLELWSMTSDIFFDNFIICADRRIVD
DWANDGWGLKKAADGAAEPGVVGQMIEAAEERPWLWVVYILTVALPVFLVILFCCSGKKQTSGM
EYKKTDAPQPDVKEEEEEKEEEKDKGDEEEEGEEKLEEKQKSDAEEDGGTVSQEEEDRKPKAET
RS**GY**CLDLKTQVQTPQGMKEISNIQVGDLVLSNTGYNEVLNVFPKSKKKSYKITLEDGKEIICS
EEHLFPTQTGEMNISGGLKEGMCLYVKEGGSGGGSGVSKGEELFTGVVPILVELDGDVNGHKFS
VRGEGEGDATNGKLTLKFICTTGKLPVPWPTLVTTFGYGVACFSRYPDHMKQHDFFKSAMPEGY
VQERTISFKDDGTYKTRAEVKFEGDTLVNRIELKGIDFKEDGNILGHKLEYNFNSHYVYITADK
QKNCIKANFKIRHNVEDGSVQLADHYQQNTPIGDGPVLLPDNHYLSHQSKLSKDPNEKRDHMVL
LEFVTAAGITHGMDELYKEQKLISEEDLGSGGSGGIKIATRKYLGKQNVYDIGVERCHNFALKN
GFIASN**CFN**EDEILNRSPRNRKPRRE*

**CANXi14- Intein Acceptor (mNG2$_{11}$, Myc)**

MEGKWLLCMLLVLGTAIVEAHDGHDDDVIDIEDDLDDVIEEVEDSKPDTTAPPSSPKVTYKAPV
PTGEVYFADSFDRGTLSGWILSKAKKDDTDDEIAKYDGKWEVEEMKESKLPGDKGLVLMSRAKH
HAISAKLNKPFLFDTKPLIVQYEVNFQNGIECGGAYVKLLSKTPELNLDQFHDKTPYTIMFGPD

KCGEDYKLHFIFRHKNPKTGIYEEKHAKRPDADLKTYFTDKKTHLYTLILNPDNSFEILVDQSV
VNSGNLLNDMTPPVNPSREIEDPEDRKPEDWDERPKIPDPEAVKPDDWDEDAPAKIPDEEATKP
EGWLDDEPEYVPDPDAEKPEDWDEDMDGEWEAPQIANPRCESAPGCGVWQRPVIDNPNYKGKWK
PPMIDNPSYQGIWKPRKIPNPDFFEDLEPFRMTPFSAIGLELWSMTSDIFFDNFIICADRRIVD
DWANDGWGLKKAADGAAEPGVVGQMIEAAEERPWLWVVYILTVALPVFLVILFCCSGKKQTSGM
EYKKTDAPQPDVKEEEEEKEEEKDKGDEEEEGEEKLEEKQKSDAEEDGGTVSQEEEDRKPKAE**T**
**RSGY**CLDLKTQVQTPQGMKEISNIQVGDLVLSNTGYNEVLNVFPKSKKKSYKITLEDGKEIICS
EEHLFPTQTGEMNISGGLKEGMCLYVKEGGSGGGSGTELNFKEWQKAFTDMMEQKLISEEDLGS
GGSGGIKIATRKYLGKQNVYDIGVERCHNFALKNGFIASN**CFN**EDEILNRSPRNRKPRRE*

**ACTN1i18 -Intein Acceptor ($mNG2_{11}$, Myc)**
MDHYDSQQTNDYMQPEEDWDRDLLLDPAWEKQQRKTFTAWCNSHLRKAGTQIENIEEDFRDGLK
LMLLLEVISGERLAKPERGKMRVHKISNVNKALDFIASKGVKLVSIGAEEIVDGNVKMTLGMIW
TIILRFAIQDISVEETSAKEGLLLWCQRKTAPYKNVNIQNFHISWKDGLGFCALIHRHRPELID
YGKLRKDDPLTNLNTAFDVAEKYLDIPKMLDAEDIVGTARPDEKAIMTYVSSFYHAFSGAQKAE
TAANRICKVLAVNQENEQLMEDYEKLASDLLEWIRRTIPWLENRVPENTMHAMQQKLEDFRDYR
RLHKPPKVQEKCQLEINFNTLQTKLRLSNRPAFMPSEGRMVSDINNAWGCLEQVEKGYEEWLLN
EIRRLERLDHLAEKFRQKASIHEAWTDGKEAMLRQKDYETATLSEIKALLKKHEAFESDLAAHQ
DRVEQIAAIAQELNELDYYDSPSVNARCQKICDQWDNLGALTQKRREALERTEKLLETIDQLYL
EYAKRAAPFNNWMEGAMEDLQDTFIVHTIEEIQGLTTAHEQFKATLPDADKERLAILGIHNEVS
KIVQTYHVNMAGTNPYTTITPQEINGKWDHVRQLVPRRDQALTEEHARQQHNERLRKQFGAQAN
VIGPWIQTKMEEIGRISIEMHGTLEDQLSHLRQYEKSIVNYKPKIDQLEGDHQLIQEALIFDNK
HTNYTMEHIRVGWEQLLTTIARTINEVENQILTRDAKGISQEQMNEFRASFNHFDR**TRSGY**CLD
LKTQVQTPQGMKEISNIQVGDLVLSNTGYNEVLNVFPKSKKKSYKITLEDGKEIICSEEHLFPT
QTGEMNISGGLKEGMCLYVKEGGSGGGSGTELNFKEWQKAFTDMMEQKLISEEDLGSGGSGGIK
IATRKYLGKQNVYDIGVERCHNFALKNGFIASN**CFN**DHSGTGPEEFKACLISLGYDIGNDPQKK
TGMMDTDDFRACLISMGYNMGEAEFARIMSIVDPNRLGVVTFQAFIDFMSRETADTDTADQVMA
SFKILAGDKNYITMDELRRELPPDQAEYCIARMAPYTGPDSVPGALDYMSFSTALYGESDL*

---

**Protein sequences for the edited proteins resulting from splicing**

Key:
Peptide edited into protein
**Bold = residues inserted for extein**
V5
FLAG

**CANX-HA (endogenous)**
MEGKWLLCMLLVLGTAIVEAHDGHDDDVIDIEDDLDDVIEEVEDSKPDTTAPPSSPKVTYKAPV
PTGEVYFADSFDRGTLSGWILSKAKKDDTDDEIAKYDGKWEVEEMKESKLPGDKGLVLMSRAKH
HAISAKLNKPFLFDTKPLIVQYEVNFQNGIECGGAYVKLLSKTPELNLDQFHDKTPYTIMFGPD
KCGEDYKLHFIFRHKNPKTGIYEEKHAKRPDADLKTYFTDKKTHLYTLILNPDNSFEILVDQSV
VNSGNLLNDMTPPVNPSREIEDPEDRKPEDWDERPKIPDPEAVKPDDWDEDAPAKIPDEEATKP
EGWLDDEPEYVPDPDAEKPEDWDEDMDGEWEAPQIANPRCESAPGCGVWQRPVIDNPNYKGKWK
PPMIDNPSYQGIWKPRKIPNPDFFEDLEPFRMTPFSAIGLELWSMTSDIFFDNFIICADRRIVD
DWANDGWGLKKAADGAAEPGVVGQMIEAAEERPWLWVVYILTVALPVFLVILFCCSGKKQTSGM

EYKKTDAPQPDVKEEEEEKEEEKDKGDEEEEGEEKLEEKQKSDAEEDGGTVSQEEEDRKPKAE**T**
**RSGY**SSSYPYDVPDYAAEY**CFN**EDEILNRSPRNRKPRRE*

**CANX-pAzF-labeled (endogenous)**
MEGKWLLCMLLVLGTAIVEAHDGHDDDVIDIEDDLDDVIEEVEDSKPDTTAPPSSPKVTYKAPV
PTGEVYFADSFDRGTLSGWILSKAKKDDTDDEIAKYDGKWEVEEMKESKLPGDKGLVLMSRAKH
HAISAKLNKPFLFDTKPLIVQYEVNFQNGIECGGAYVKLLSKTPELNLDQFHDKTPYTIMFGPD
KCGEDYKLHFIFRHKNPKTGIYEEKHAKRPDADLKTYFTDKKTHLYTLILNPDNSFEILVDQSV
VNSGNLLNDMTPPVNPSREIEDPEDRKPEDWDERPKIPDPEAVKPDDWDEDAPAKIPDEEATKP
EGWLDDEPEYVPDPDAEKPEDWDEDMDGEWEAPQIANPRCESAPGCGVWQRPVIDNPNYKGKWK
PPMIDNPSYQGIWKPRKIPNPDFFEDLEPFRMTPFSAIGLELWSMTSDIFFDNFIICADRRIVD
DWANDGWGLKKAADGAAEPGVVGQMIEAAEERPWLWVVYILTVALPVFLVILFCCSGKKQTSGM
EYKKTDAPQPDVKEEEEEKEEEKDKGDEEEEGEEKLEEKQKSDAEEDGGTVSQEEEDRKPKAE**T**
**RSGY**SSS(pAzF)AEY**CFN**EDEILNRSPRNRKPRRE*

**α-actinin-1 – HA (endogenous)**
MDHYDSQQTNDYMQPEEDWDRDLLLDPAWEKQQRKTFTAWCNSHLRKAGTQIENIEEDFRDGLK
LMLLLEVISGERLAKPERGKMRVHKISNVNKALDFIASKGVKLVSIGAEEIVDGNVKMTLGMIW
TIILRFAIQDISVEETSAKEGLLLWCQRKTAPYKNVNIQNFHISWKDGLGFCALIHRHRPELID
YGKLRKDDPLTNLNTAFDVAEKYLDIPKMLDAEDIVGTARPDEKAIMTYVSSFYHAFSGAQKAE
TAANRICKVLAVNQENEQLMEDYEKLASDLLEWIRRTIPWLENRVPENTMHAMQQKLEDFRDYR
RLHKPPKVQEKCQLEINFNTLQTKLRLSNRPAFMPSEGRMVSDINNAWGCLEQVEKGYEEWLLN
EIRRLERLDHLAEKFRQKASIHEAWTDGKEAMLRQKDYETATLSEIKALLKKHEAFESDLAAHQ
DRVEQIAAIAQELNELDYYDSPSVNARCQKICDQWDNLGALTQKRREALERTEKLLETIDQLYL
EYAKRAAPFNNWMEGAMEDLQDTFIVHTIEEIQGLTTAHEQFKATLPDADKERLAILGIHNEVS
KIVQTYHVNMAGTNPYTTITPQEINGKWDHVRQLVPRRDQALTEEHARQQHNERLRKQFGAQAN
VIGPWIQTKMEEIGRISIEMHGTLEDQLSHLRQYEKSIVNYKPKIDQLEGDHQLIQEALIFDNK
HTNYTMEHIRVGWEQLLTTIARTINEVENQILTRDAKGISQEQMNEFRASFNHFDR**TRSGY**SSS
YPYDVPDYAAEY**CFN**DHSGTGPEEFKACLISLGYDIGNDPQKKTGMMDTDDFRACLISMGYNMG
EAEFARIMSIVDPNRLGVVTFQAFIDFMSRETADTDTADQVMASFKILAGDKNYITMDELRREL
PPDQAEYCIARMAPYTGPDSVPGALDYMSFSTALYGESDL*

**V5-CANX-HA-FLAG**
MGKPIPNPLLGLDSTGGSGGMEGKWLLCMLLVLGTAIVEAHDGHDDDVIDIEDDLDDVIEEVED
SKPDTTAPPSSPKVTYKAPVPTGEVYFADSFDRGTLSGWILSKAKKDDTDDEIAKYDGKWEVEE
MKESKLPGDKGLVLMSRAKHHAISAKLNKPFLFDTKPLIVQYEVNFQNGIECGGAYVKLLSKTP
ELNLDQFHDKTPYTIMFGPDKCGEDYKLHFIFRHKNPKTGIYEEKHAKRPDADLKTYFTDKKTH
LYTLILNPDNSFEILVDQSVVNSGNLLNDMTPPVNPSREIEDPEDRKPEDWDERPKIPDPEAVK
PDDWDEDAPAKIPDEEATKPEGWLDDEPEYVPDPDAEKPEDWDEDMDGEWEAPQIANPRCESAP
GCGVWQRPVIDNPNYKGKWKPPMIDNPSYQGIWKPRKIPNPDFFEDLEPFRMTPFSAIGLELWS
MTSDIFFDNFIICADRRIVDDWANDGWGLKKAADGAAEPGVVGQMIEAAEERPWLWVVYILTVA
LPVFLVILFCCSGKKQTSGMEYKKTDAPQPDVKEEEEEKEEEKDKGDEEEEGEEKLEEKQKSDA
EEDGGTVSQEEEDRKPKAETRS**GY**SSSYPYDVPDYAAEY**CFN**EDEILNRSPRNRKPRREGGSGG
DYKDDDDK*

**ACTB-HA-FLAG (i2)**

MDDDIAALVVDNGSGMCKAGFAGDDAPRAVFPSIVGRPRHQ**GY**SSSYPYDVPDYAAEY**CFN**GVM
VGMGQKDSYVGDEAQSKRGILTLKYPIEHGIVTNWDDMEKIWHHTFYNELRVAPEEHPVLLTEA
PLNPKANREKMTQIMFETFNTPAMYVAIQAVLSLYASGRTTGIVMDSGDGVTHTVPIYEGYALP
HAILRLDLAGRDLTDYLMKILTERGYSFTTTAEREIVRDIKEKLCYVALDFEQEMATAASSSSL
EKSYELPDGQVITIGNERFRCPEALFQPSFLGMESCGIHETTFNSIMKCDVDIRKDLYANTVLS
GGTTMYPGIADRMQKEITALAPSTMKIKIIAPPERKYSVWIGGSILASLSTFQQMWISKQEYDE
SGPSIVHRKCFGGSGGDYKDDDDK*

**ACTB-HA-FLAG (i5)**
MDDDIAALVVDNGSGMCKAGFAGDDAPRAVFPSIVGRPRHQGVMVGMGQKDSYVGDEAQSKRGI
LTLKYPIEHGIVTNWDDMEKIWHHTFYNELRVAPEEHPVLLTEAPLNPKANREKMTQIMFETFN
TPAMYVAIQAVLSLYASGRTTGIVMDSGDGVTHTVPIYEGYALPHAILRLDLAGRDLTDYLMKI
LTERGYSFTTTAEREIVRDIKEKLCYVALDFEQEMATAASSSSLEKSYELPDGQVITIGNERFR
CPEALFQPSFLGMESCGIHETTFNSIMKCDVDIRKDLYANTVLSGGTTMYPGIADRMQKEITAL
APSTMKIK**GY**SSSYPYDVPDYAAEY**CFN**IIAPPERKYSVWIGGSILASLSTFQQMWISKQEYDE
SGPSIVHRKCFGGSGGDYKDDDDK*

**Chk1-FLAG (P350F, D351N)**
MGDYKDDDDKMAVPFVEDWDLVQTLGEGAYGEVQLAVNRVTEEAVAVKIVDMKRAVDCPENIKK
EICINKMLNHENVVKFYGHRREGNIQYLFLEYCSGGELFDRIEPDIGMPEPDAQRFFHQLMAGV
VYLHGIGITHRDIKPENLLLDERDNLKISDFGLATVFRYNNRERLLNKMCGTLPYVAPELLKRR
EFHAEPVDVWSCGIVLTAMLAGELPWDQPSDSCQEYSDWKEKKTYLNPWKKIDSAPLALLHKIL
VENPSARITIPDIKKDRWYNKPLKKGAKRPRVTSGGVSESPSGFSKHIQSNLDFSPVNSASSEE
NVKYSSSQPEPRTGLSLWDTSPSYIDKLVQGISFSQPTCFNHMLLNSQLLGTPGSSQNPWQRLV
KRMTRFFTKLDADKSYQCLKETCEKLGYQWKKSCMNQVTISTTDRRNNKLIFKVNLLEMDDKIL
VDFRLSKGDGLEFKRHFLKIKGKLIDIVSSQKVWLPAT*

**c-Myc-FLAG (A44G,P45Y, S71F, P72N)**
MPLNVSFTNRNYDLDYDSVQPYFYCDEEENFYQQQQQSELQPPGYSEDIWKKFELLPTPPLSPS
RRSGLCFNSYVAVTPFSLRGDNDGGGGSFSTADQLEMVTELLGGDMVNQSFICDPDDETFIKNI
IIQDCMWSGFSAAAKLVSEKLASYQAARKDSGSPNPARGHSVCSTSSLYLQDLSAAASECIDPS
VVFPYPLNDSSSPKSCASQDSSAFSPSSDSLLSSTESSPQGSPEPLVLHEETPPTTSSDSEEEQ
EDEEEIDVVSVEKRQAPGKRSESGSPSAGGHSKPPHSPLVLKRCHVSTHQHNYAAPPSTRKDYP
AAKRVKLDSVRVLRQISNNRKCTSPRSSDTEENVKRRTHNVLERQRRNELKRSFFALRDQIPEL
ENNEKAPKVVILKKATAYILSVQAEEQKLISEEDLLRKRREQLKHKLEQLRNSCAGSGDYKDDD
DK*
